# Behavioral, physiological, and neural signatures of surprise during naturalistic sports viewing

**DOI:** 10.1101/2020.03.26.008714

**Authors:** James W. Antony, Thomas H. Hartshorne, Ken Pomeroy, Todd M. Gureckis, Uri Hasson, Samuel D. McDougle, Kenneth A. Norman

## Abstract

Surprise signals a discrepancy between past and current beliefs. It is theorized to be linked to affective experiences, the creation of particularly resilient memories, and segmentation of the flow of experience into discrete perceived events. However, the ability to precisely measure naturalistic surprise has remained elusive. We used advanced basketball analytics to derive a quantitative measure of surprise and characterized its behavioral, physiological, and neural correlates in human subjects observing basketball games. We found that surprise was associated with segmentation of ongoing experiences, as reflected by subjectively perceived event boundaries and shifts in neocortical patterns underlying belief states. Interestingly, these effects differed by whether surprising moments contradicted or bolstered current predominant beliefs. Surprise also positively correlated with pupil dilation, activation in subcortical regions associated with dopamine, game enjoyment, and long-term memory. These investigations support key predictions from event segmentation theory and extend theoretical conceptualizations of surprise to real-world contexts.

## Introduction

As events in the world unfold, the brain rapidly adjusts its predictions of what will happen next. Of course, our predictions are not always correct – when they are inaccurate, we often experience surprise (i.e., unsigned prediction error) (Jang et al., 2019; Kutas and Hillyard, 1984; O’Reilly et al., 2013; Rouhani et al., 2018, 2020). Surprise is theorized to be critical for learning and memory (Rescorla and Wagner, 1972; Sinclair and Barense, 2018), updating our beliefs about the structure of the world (Sutton and Barto, 1998), and demarcating events in the continuous flow of time (Franklin et al., 2020). Moreover, although people typically prefer certainty about outcomes that are instrumental for survival (Bromberg-Martin and Hikosaka, 2009), in domains with non-instrumental information, such as narratives, music, and sports, people tend to prefer violations of their expectations (Ely et al., 2015; Gan et al., 1997; Gold et al., 2019), suggesting that surprise is often a rewarding affective experience.

Although surprise has been elegantly operationalized in laboratory experiments, in more naturalistic settings it is difficult to precisely characterize. Most experiments measure surprise in the context of discrete temporal units (e.g., trials) involving repeated sensory cues, rather than as a probabilistic belief state that is continuously updated over time (Hutchinson and Barrett, 2019). Here, we leveraged naturalistic stimuli (video clips of basketball games) in which we could quantify how people continuously update their predictions about an outcome (which team will win).

First, we describe and validate our model of surprise. Then, we show how surprise relates to perceived event boundaries, the segmentation of neural event states, and pupil dilation. Finally, we show that surprise predicts both subjective enjoyment and neural signatures of reward and leads to improved long-term memory for events.

## Results

### Calculating and validating surprise during basketball viewing

Self-ascribed basketball fans (*N* = 20, 6 female) underwent eye tracking and fMRI scans while watching the final five minutes of nine games from the 2012 men’s NCAA® college basketball (“March Madness”) tournament (Fig 1A). Subjects reported their preferred team (if any) and enjoyment after each game and freely recalled the games from memory after each set of three. We operationalized predictions using a “win probability” model trained on a corpus of games from the 2012 regular season that was based on four factors: the difference in score between two teams (oriented as positive when the “home” team was winning and negative when they were losing), the amount of time remaining, which team was in possession of the ball, and the relative strength of the two teams. The model provides the win probability updated for each possession based on similar game states in the corpus (Fig 1B; Fig S1). Score difference is the most important factor. Score difference and the team possessing the ball become more important as time elapses. Relative team strength factors into the model by taking the “expected” score difference (from the public website of an expert basketball analyst, www.kenpom.com), dividing this by the total number of seconds in the game (to get an estimate of how much the stronger team was supposed to outscore the worse team, in units of “points per second”), and multiplying by the number of seconds remaining in the game; this approach assumes that the stronger team’s score advantage accrues at a constant rate (e.g., if a team is supposed to win by 20, that translates into an expected 10 point differential in each half; see Methods for more information). We validated these predictions against those of the expert basketball analyst on both a held-out subset of games from the corpus (Fig 1C, bottom) and the tournament games viewed in this experiment (Fig 1C, top).

**Fig 1.**
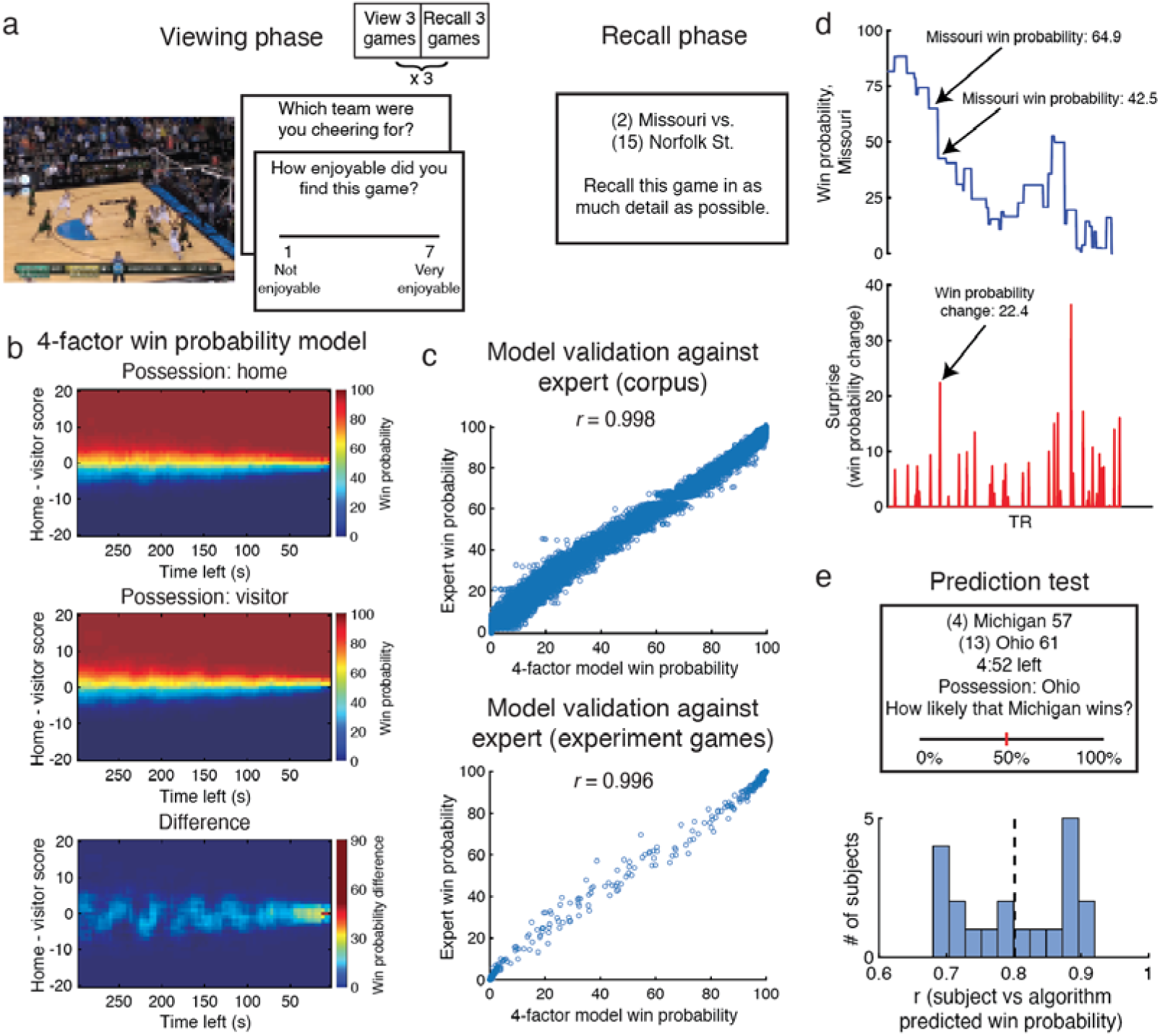
Experimental procedure and surprise calculation and validation. a) Subjects undergoing eye-tracking and fMRI viewed and recalled nine NCAA basketball games. Game screenshot is blurred here for copyright reasons. b) Model predicting win probabilities (color scale) based on the score difference (y axis), time remaining (x axis), team in possession of the ball, and relative strength of the teams (not shown here, see text). Probabilities are plotted separately for when the home team possesses the ball (top) and when the visiting team possesses the ball (middle), with the difference between these scenarios on the bottom. c) Model predictions for each possession corresponded tightly with those from an expert sports analyst in corpus data (top) and in games used in this experiment (bottom). d) (top) Win probabilities from the model chart the likelihood of one team winning across time points in the scanner (TRs). (bottom) Surprise was defined as the unsigned change in this time course. e) (top) Subject predictions were tested on scenarios from different games given at the end of the experiment. (bottom) Histogram showing that subject predictions were correlated highly with predictions from the model. The vertical dashed line indicates the mean.

One can think of each subject’s brain state over the course of a game as traversing a state space of predictions about the likely winner (Fig 1C,D). Our win probability measure was computed so that it updates after each possession change (i.e., after scores or turnovers – whenever the other team obtains the ball). Consequently, we computed surprise as the absolute value of the change in the win probability time course at each possession boundary (Fig 1D). Note that, in other work, the term “surprise” has been used to refer to prediction errors (differences between expected and actual moment-to-moment outcomes) that can (Faraji et al., 2018), but do not always (O’Reilly et al., 2013), lead to updates in one’s underlying beliefs about the world. For the analyses reported here, we reserve the use of the term “surprise” to refer to model-derived changes in estimated win probabilities, which are reflected in subjects’ internal estimates of win probability (as described below; we return to these points in the *Discussion*).

To validate that subjects represented these probabilities in some form, at the end of the experiment we presented them with a test featuring scenarios from different 2012 tournament games that they had not previously viewed (Fig 1E). In this test, subjects rated the likelihood that the home team would win for every possession (starting with five minutes left) when given the seeds of the teams, the score, the amount of time remaining, and the team in possession of the ball. The distribution of probabilities in this test closely resembled those from viewed games (Fig S1). Indeed, subject responses correlated highly with the win probabilities specified by our algorithm (mean *r* = 0.80, range = 0.68 to 0.91, *p* < 0.001; Fig 1E), validating its use for approximating surprise.

### Surprise correlates with subjective event segmentation, the segmentation of neural states, and pupil dilation

Event segmentation theory (EST) posits that humans naturally segment their ongoing stream of experience and create internal event models to predict upcoming events (Zacks et al., 2007, 2011). Violations of these predictions (i.e., surprises) are thought to coincide with event boundaries, which are reflected in subjective segmentations of continuous perception (Zacks et al., 2011), pronounced shifts in neural states (Baldassano et al., 2017), and physiological changes such as pupil dilations (Braem et al., 2015; Clewett et al., 2019a; Filipowicz et al., 2020; Preuschoff et al., 2011; Yu and Dayan, 2005). Here, we investigated how surprise relates to these three measurements.

First, we wanted to examine how a separate group of basketball fans perceived game events and how these judgments related to surprise. These fans (*N* = 15, 8 female) watched the games outside of the scanner and responded when they perceived the ending and new beginning of “game units” at the coarsest level that was meaningful to them (Newtson, 1973) (see *Methods* for instructions). Overall, subjects responded anywhere from 5 to 150 times (66.1 ± 11.7) across the 9 games (Fig 2A). Based on these responses, we computed a *subjective boundary agreement* score for each possession boundary (e.g., score or change in possession), quantifying the agreement across subjects (ranging from 0 for no agreement to 1 for perfect agreement) that a game unit ended within 2 s of that possession boundary. At each possession boundary (157 total, across all 9 games), we therefore had a measure of subjective boundary agreement, which we then correlated (across the 157 possession boundaries) with model-derived surprise. We tested this correlation against a null distribution of permutations, where we circularly shifted surprise values across possessions within each game and then concatenated all 157 values together for each permutation. Partially supporting the predictions from EST outlined above, subjective boundary agreement showed a marginal correlation with surprise (*r* = 0.29, *p* = 0.07) (Fig 2C).

**Fig 2:**
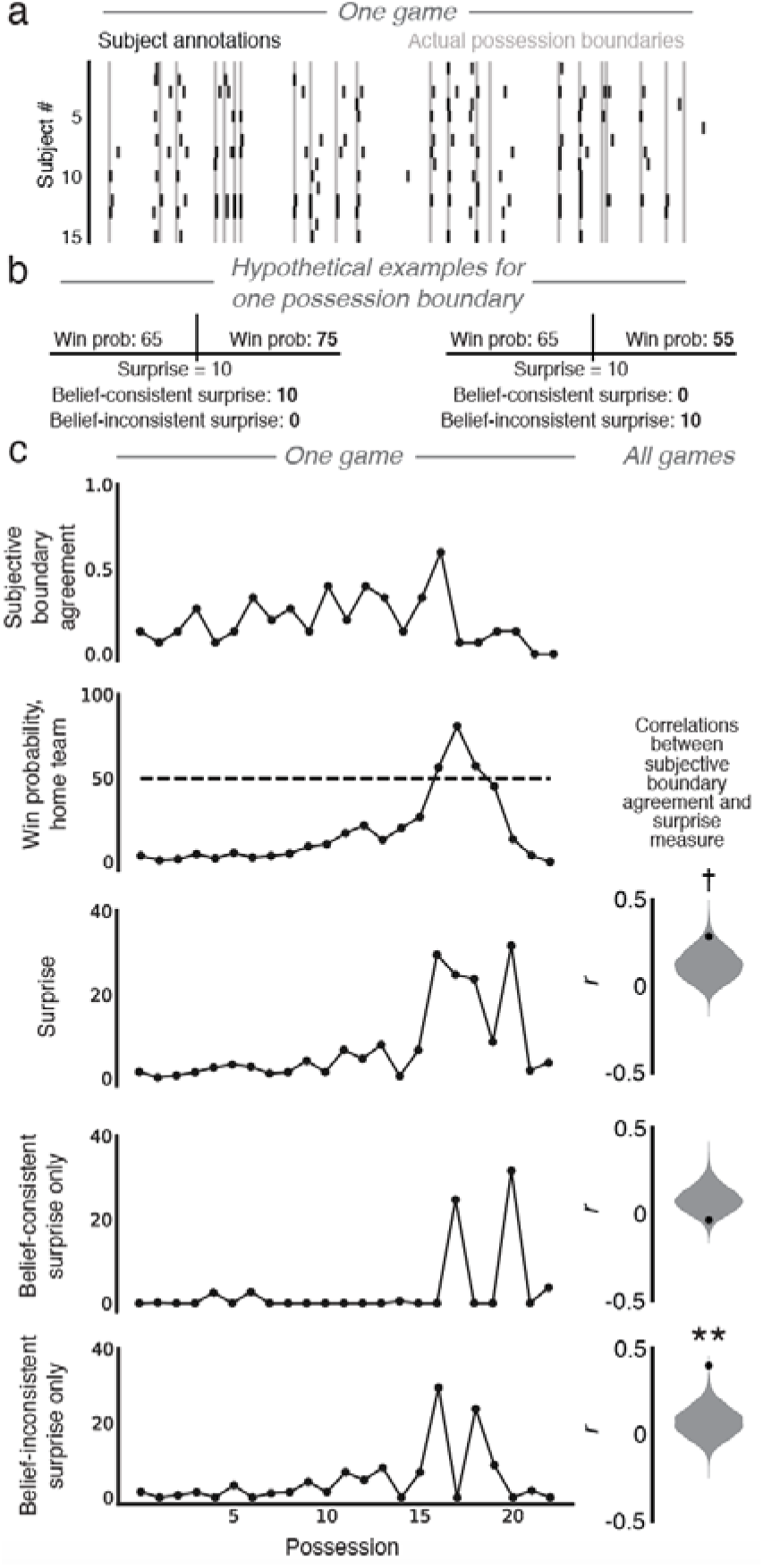
Perceived event boundaries correlate with surprise measures. a) Each subject’s annotations demarcating game units are shown in rows (black ticks) amidst actual possession boundaries for one game (gray); note that subjects making the annotations were from a separate cohort. (b) Hypothetical examples show how belief-consistent surprise and belief-inconsistent surprise were calculated. (c) (left) Subjective boundary agreement (the proportion of subjects who marked an event boundary at each possession boundary) is plotted for one game against win probability for the home team, surprise, belief-consistent surprise, and belief-inconsistent surprise at those boundaries. (right) Subjective boundary agreement was marginally correlated with surprise and significantly correlated with belief-inconsistent surprise. Black dots indicate true values versus smoothed null distributions in grey. †: 0.05 < *p* < 0.10. **: *p* < 0.01.

We next intuited that, while subjects can represent these predictions in a graded fashion (Fig 1C), they could also hold a binary belief about which team is more likely to win at a particular moment (e.g., team X currently has > 50% chance of winning) (Johnson et al., 2020). If so, there could be a qualitative difference in surprise based on whether new evidence is consistent with this belief (e.g., team X scores, furthering their lead) or inconsistent with this belief (e.g., team Y scores, cutting into team X’s lead). To explore this, we created the constructs of belief-consistent surprise and belief-inconsistent surprise (Fig 2B,C). Belief-consistent surprise is equivalent to surprise if the change in win probability involves the team with the higher win probability becoming even more likely to win; it is zero otherwise. Belief-inconsistent surprise is equivalent to surprise if the change in win probability involves the team with the higher win probability becoming less likely to win; it is zero otherwise. We hypothesized that belief-inconsistent surprise would entail more updating of one’s event model (i.e., one’s understanding of what is happening), and thus – according to Event Segmentation Theory (Zacks et al., 2007, 2011) – would be more likely to trigger segmentation (Fig 2B). To test this, we next correlated subjective boundary agreement across possession boundaries separately with belief-consistent and belief-inconsistent surprise (Fig 2C). The correlation was significant for belief-inconsistent but not belief-consistent surprise, and the two significantly differed (inconsistent: *r* = 0.40, *p* < 0.001; consistent: *r* = −0.03, *p* = 0.10; difference: *r* = 0.43, *p* < 0.001, permutation tests circularly shuffling possessions within each game). Taken together, these results suggest that event segmentation marginally increases with surprise and, rather than simply demarcating bidirectional probabilistic changes in belief, segmentation is especially robust when new information conflicts with the predominant belief.

We next attempted to capture signatures of the segmentation of continuous experience in neural data, leveraging Hidden Markov models (HMMs) to analyze BOLD responses in fMRI (Baldassano et al., 2017; Chang et al., 2019). HMMs are data-driven algorithms that probabilistically segment data into stable states and discrete shifts between those states. We used a specialized HMM variant developed by Baldassano et al. (2017) that is optimized for event segmentation (i.e., identifying “jumps” in neural patterns). This HMM variant assumes that the neural time series can be modeled by transitioning through some number of discrete states, never returning to previous states (Baldassano et al., 2017). Here we used HMMs to address three questions: First, do HMM state transitions naturally align with actual game possession boundaries? Second, following from the hypothesis that surprise leads to event segmentation, does surprise predict more frequent HMM state transitions across possessions and does greater cumulative surprise lead to more neural event states across games? Finally, are HMM state transitions more likely to occur at moments of belief-inconsistent or belief-consistent surprise? We addressed these questions using neural activity from primary visual cortex (V1), precuneus, and medial prefrontal cortex (mPFC) as *a priori* regions-of-interest (ROIs) (Fig 3A). We predicted that V1 would mainly track small-time-scale changes with salient sensory features (Baldassano et al., 2017; Lerner et al., 2011), like possession changes, but could also be modulated by top-down processes (Hindy et al., 2016; Hutchinson and Barrett, 2019). For precuneus, a node in the default mode network, we predicted that state changes would occur on a longer time-scale and reflect subjective segmentations (as was found previously in Baldassano et al., 2017), and that these state changes would occur more frequently with higher surprise. Finally, we predicted that representations in mPFC, a higher-level region involved in abstract inference and the representation of abstract states (Hampton et al., 2006; Starkweather et al., 2018; Takahashi et al., 2011; Wilson et al., 2014), would exhibit state changes on the longest time-scale, and would also occur more frequently with higher surprise, reflecting shifts in the broader narrative of the game. We also predicted that correlations between neural state changes in higher-order areas (precuneus, mPFC) and surprise would be largest when we focus on *belief-inconsistent* (vs. belief-consistent) surprise, for the reasons noted above: Belief-inconsistent surprise entails a greater update to one’s event model and thus should be more likely to trigger segmentation.

**Fig 3:**
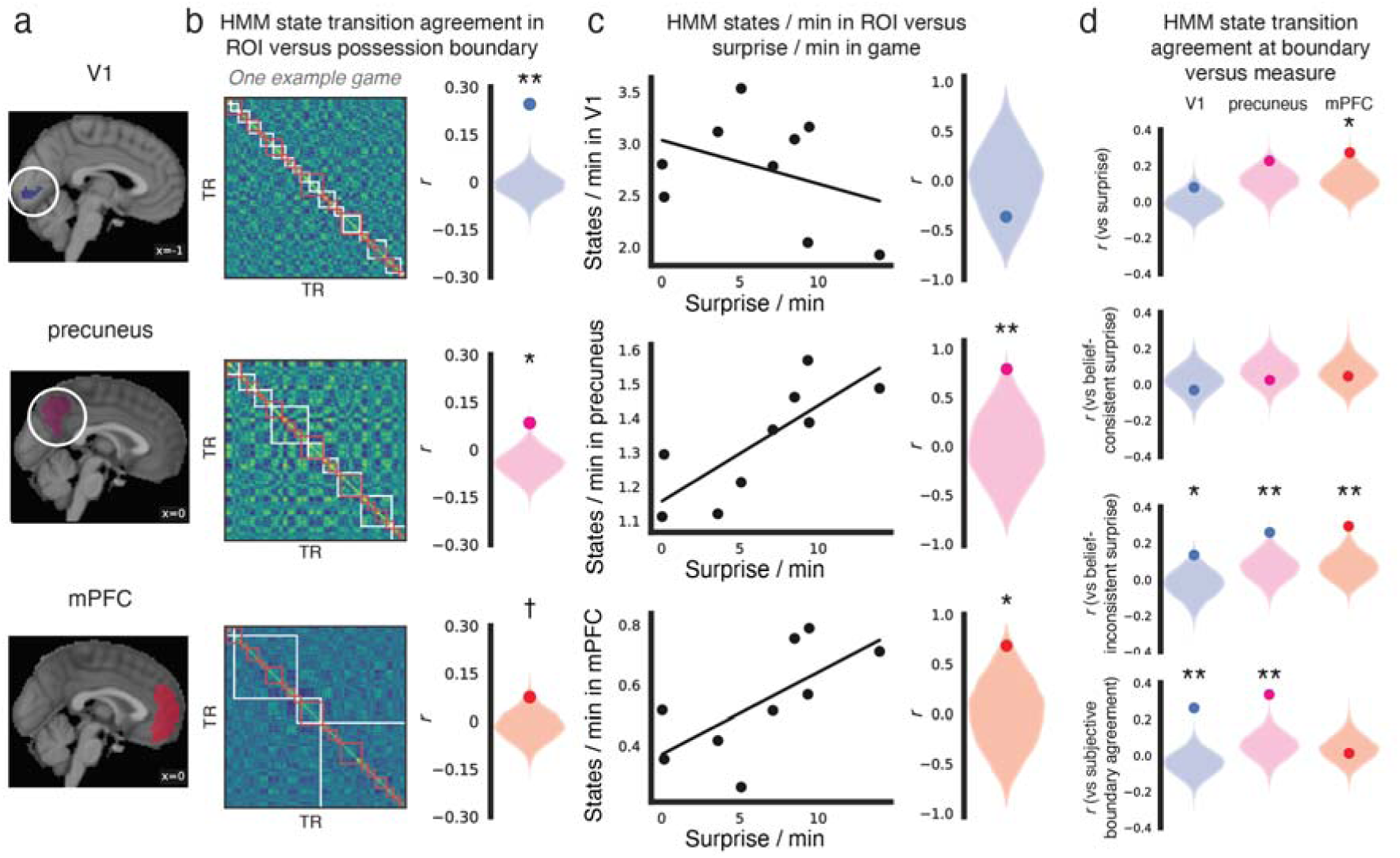
Surprise predicts neural event boundaries. a) V1 (top), precuneus (middle), and mPFC (bottom) ROIs are shown. b) (left) For each ROI, the figure shows representative correlation matrices (for a single game and single subject) indicating the similarity between spatial patterns of activity in that ROI for every pair of TRs (the diagonal is trivially 1 as it plots the similarity of each TR to itself). White boxes show groups of TRs that were identified by our Hidden Markov Model (HMM) analysis as belonging to the same event. Actual possession boundaries are shown in red. (right) Correlations between the time series of HMM state transitions and the time series of true possession boundaries are plotted (dark circles), compared to null distributions. This relationship was highest in V1. C) Correlations across games (dots) show the number of states per minute in each region versus surprise per minute (left), which was significant in mPFC and precuneus (right). d) Across possession boundaries, HMM state transition agreement (the proportion of subjects showing an HMM state transition in the time window around the boundary) was correlated against surprise, belief-consistent surprise, belief-inconsistent surprise, and subjective boundary agreement for each ROI, compared to null distributions. Surprise was significantly (positively) correlated with HMM state transition agreement in mPFC. Belief-inconsistent (but not belief-consistent) surprise was also significantly correlated with HMM state transition agreement in V1, precuneus, and mPFC. Subjective boundary agreement correlated significantly with HMM state transition agreement in V1 and precuneus. †: 0.05 *p* < 0.10. *: *p* < 0.05. **: *p* < 0.01.

To address the first question (i.e., do HMM state transitions naturally align with actual game possession boundaries), we first calculated (for each region of interest, and for each time point) HMM state transition agreement, operationalized as a single value representing the proportion of subjects showing at least one HMM state transition within a 15-s window (±7 s) of each time point. We then correlated this across-subject measure with the true time course of possession changes, and we compared this to two distinct kinds of null distributions: one where we shuffled possession order within games (preserving possession length) and one where we circularly shifted time courses within games (preserving the temporally autocorrelated structure of possession boundaries). In agreement with our prediction, the time course of neural state transitions in V1 was significantly correlated with the time course of ground-truth possession changes, which involve substantial changes in visual features (Fig 3B; *r* = 0.25, *p* < 0.001, via permutation tests shuffling possession order; *p* < 0.001, via permutation tests with circular shifts). This effect was significant in precuneus (*r* = 0.084, *p* = 0.003 by shuffling possessions, *p* = 0.04 by circular shift) and marginally significant in mPFC (*r* = 0.075, *p* = 0.06 by shuffling possessions, *p* = 0.07 by circular shift); the effect was also significantly weaker in those regions than in V1 (*r* difference, V1 – precuneus: *r* = 0.16, *p* = 0.04; V1 – mPFC: *r* = 0.17, *p* = 0.007).

Next, we asked whether surprise is associated with a greater number of neural event states, both across games and across possessions. Across games, we correlated the mean surprise per minute in each game and the cross-validated best-fitting number of states per minute (averaged across subjects) within each ROI. This relationship was significant in precuneus (*r* = 0.79, *p* = 0.007) and mPFC (*r* = 0.68, *p* = 0.04, via permutation tests that shuffle games), but in V1 this was not significant (*r* = −0.37, *p* = 0.34) and was significantly lower than in mPFC (*r* difference = 1.05, *p* = 0.03) and precuneus (*r* difference = 1.16, *p* = 0.015) (Fig 3C). Observed representational shifts did not correlate with potential confounding factors such as the number of possessions per minute (precuneus: *r* = −0.38, *p* = 0.31; mPFC: *r* = −0.23, *p* = 0.53), nor the total amount of visual motion per minute (precuneus: *r* = 0.15, *p* = 0.70; mPFC: *r* = 0.24, *p* = 0.53). Moreover, the relationship with mean surprise per minute remained significant or marginally significant in a regression model controlling for these factors in precuneus (*p* = 0.03, via permutation tests that shuffle games) and mPFC (*p* = 0.07).

To investigate the effects of surprise at the finer temporal resolution of possessions, we calculated HMM state transition agreement for each possession boundary, operationalized as the proportion of subjects showing at least one HMM state transition in the 15-s window spanning the possession boundary (±7 s around the boundary). At each possession boundary (157 total, across all 9 games), we therefore had a measure of HMM state transition agreement, which we correlated (across the 157 possession boundaries) with model-derived surprise; we tested this correlation against a null distribution of permutations, where we circularly shifted surprise values across possessions within each game before concatenating them together. In contrast to the first analysis in this section, which looked at whether HMM state transitions were more likely to occur at possession boundaries (vs. other time points), this analysis “zooms in” on possession boundaries and asks whether the occurrence of HMM state transitions at these time points is modulated by surprise. We found that the correlation between HMM state transition agreement at possession boundaries and surprise was significant in mPFC (*r* = 0.27, *p* = 0.028), but this relationship was not significant in V1 (*r* = 0.08, *p* = 0.19) nor in precuneus (*r* = 0.23, *p* = 0.12) (Fig 3D) (all *r* differences between ROIs, *p* > 0.4). Therefore, mPFC pattern shifts occur more frequently with greater surprise.

As noted above, we also predicted that (across possession boundaries) HMM state transition agreement in mPFC and precuneus would correlate better with belief-inconsistent surprise than belief-consistent surprise. Consistent with this prediction, correlations were significant for belief-inconsistent surprise in all three regions (precuneus: *r* = 0.26, *p* = 0.008; mPFC: *r* = 0.29, *p* = 0.004; V1: *r* = 0.13, *p* = 0.03; all *r* differences between ROIs, *p* > 0.47). None of the regions showed a significant correlation for belief-consistent surprise (V1: *r* = −0.03, *p* = 0.49; precuneus: *r* = 0.025, *p* = 0.51; mPFC: *r* = 0.046, *p* = 0.83; all *r* differences between ROIs, *p* > 0.72), and there was a significant difference between the correlations for belief-inconsistent and belief-consistent surprise in precuneus (*r* = 0.23, *p* = 0.047) and mPFC (*r* = 0.25, *p* = 0.047), but not V1 (*r* = 0.16, *p* = 0.11) (all *r* differences between ROIs for this measure, *p* > 0.77) (Fig 3D). Taken together, these results accord with neural predictions of EST (Zacks et al., 2007, 2011), specifically that surprising outcomes – especially those that counter one’s current belief and thereby increase uncertainty – coincide with “resetting” neural representations in higher-level brain regions (e.g., precuneus, mPFC) that are involved in event processing (Nassar et al., 2019; Shin and Dubrow, 2020).

Finally, we asked whether HMM state transition agreement correlated with subjective boundary agreement (computed in our non-fMRI subjects) concatenated across 157 possessions, given that both measures correlated with surprise (and specifically belief-inconsistent surprise). We found that HMM state transition agreement in precuneus (*r* = 0.34, *p* < 0.001) and V1 (*r* = 0.26, *p* < 0.001) correlated with subjective boundary agreement, whereas HMM state transition agreement in mPFC did not (*r* = 0.01, *p* = 0.72) (*r* difference, V1 – precuneus: *r* = −0.07, *p* = 0.75; V1 – mPFC: *r* = 0.25, *p* < 0.001; precuneus – mPFC: *r* = 0.32, *p* < 0.001).

In sum, neural states in these three regions appear to change on different time-scales, and they are most strongly modulated by different variables. V1 state changes occur rapidly and track possession boundaries significantly better than the other two regions; V1 state changes also track belief-inconsistent surprise and subjective boundary agreement. Precuneus state changes occur at a more moderate rate. Like V1, precuneus state changes track belief-inconsistent surprise and subjective boundary agreement; the latter finding conceptually replicates the results of a previous study that used a narrative movie stimulus (Baldassano et al., 2017; Zadbood et al., 2017). Lastly, mPFC changes occur sparsely (less than once per minute), but they also occur preferentially at surprising moments and more often when events conflict with the current belief; mPFC changes were not significantly related to subjective event boundaries (perhaps due to a timescale mismatch – mPFC state changes, with an average of 2.5 transitions / game, occurred much less frequently than subjective event boundaries, with an average of 7.3 / game). The overall pattern of effects in mPFC is consistent with mPFC changing its state during moments of behavioral uncertainty (Karlsson et al., 2012) or salient changes in environmental structure (D’Acremont et al., 2013; Durstewitz et al., 2010; Nassar et al., 2019).

Additionally, we performed exploratory analyses asking how HMM state transitions correspond to the above surprise measures in other parts of the neocortex by repeating these across-possession analyses in 48 bilateral cortical parcels from the Harvard-Oxford Brain Atlas (Fig S2, full details in Table S1). These results were largely consistent with those above: The paracingulate cortex parcel, which best overlaps our mPFC ROI, was among the regions showing the strongest correlations for both surprise (*r* = 0.235, *p* = 0.02, permutation test shuffling possessions; after three other parcels) and belief-inconsistent surprise (*r* = 0.26, *p* = 0.01; after seven other parcels). The precuneus parcel showed strong correlations with belief-inconsistent surprise (*r* = 0.27, *p* = 0.002; after five other parcels) and subjective boundary agreement (*r* = 0.24, *p* < 0.006; after nine other parcels).

Previous accounts have linked event segmentation and surprise with pupil dilation (Braem et al., 2015; Clewett et al., 2019a; Filipowicz et al., 2020; Preuschoff et al., 2011; Yu and Dayan, 2005), which we investigated next. There are challenges analyzing pupil dilation with video stimuli, as pupil area measurements with conventional eye trackers differ by the gaze location of the eye (Hayes and Petrov, 2017), decrease with global and local visual luminance (Page, 1941), and increase with salient sounds (Nassar et al., 2012). We addressed the first challenge by normalizing the measurements within x- and y-coordinate bins according to the gaze location (Fig 4A). We addressed the second and third challenges by including the following sensory variables (computed for each second of the game broadcast) in a linear model relating surprise to pupil dilation: global luminance of the entire video, local luminance surrounding the eye location, and the auditory envelope from the broadcast (Fig 4B). Additionally, we created regressors for the following: global and local video motion based on the change in pixel-by-pixel luminance across video frames; the fundamental frequency (f0) of the commentator’s speech to capture changes in prosody; the amount of game remaining based on the game time at the start of the possession; and the position of the ball in the telecast (right or left side of the court).

**Fig 4.**
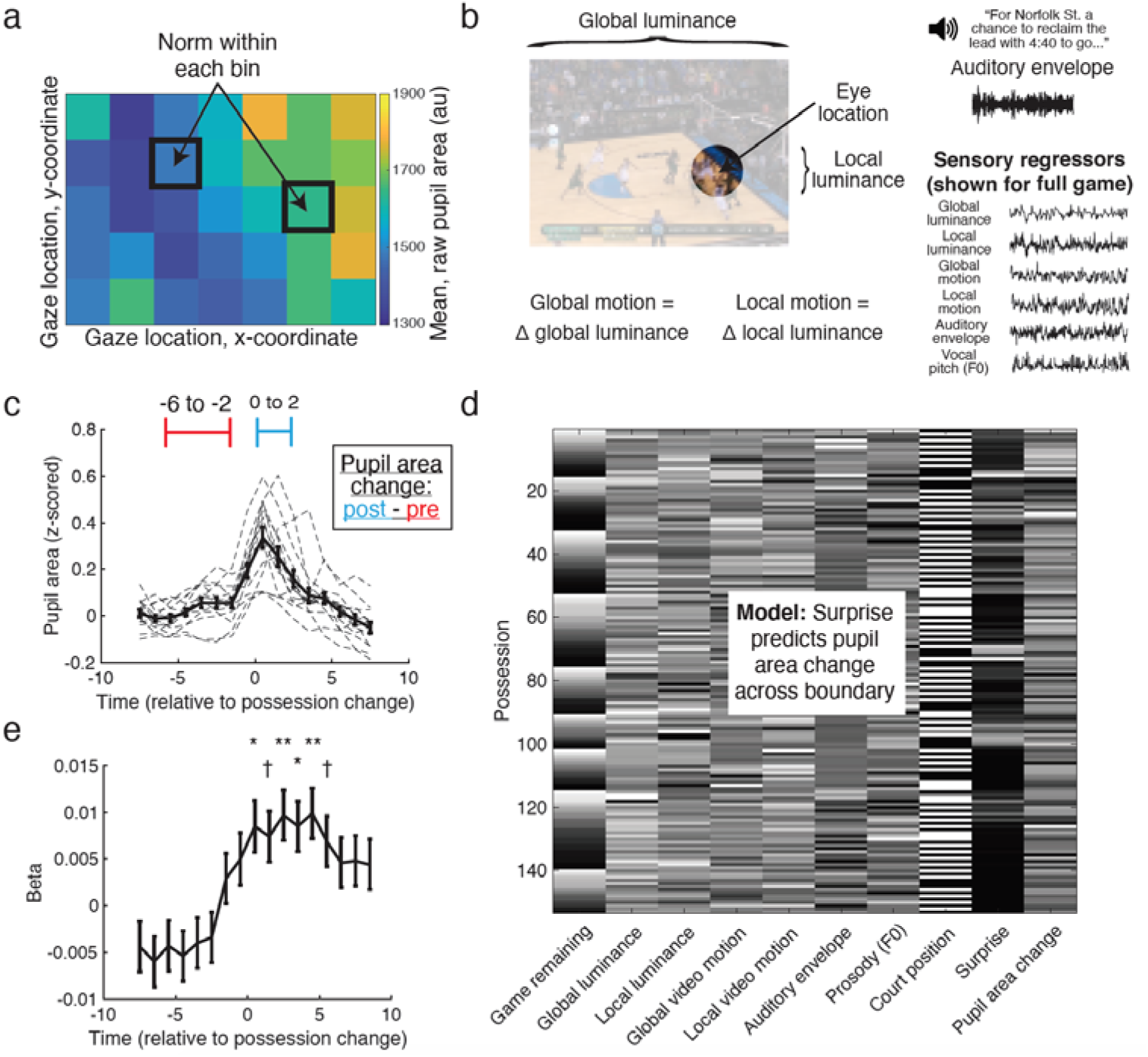
Surprise increases with pupil dilation across possession boundaries. a) Confounds due to gaze location were addressed by splitting the screen into x- and y-coordinate bins and norming the raw pupil area values within each bin. b) (top left) Luminance for each TR was calculated as an average for the entire screen (global luminance) as well as for a 2° radius around the current location of the eye (local luminance). (bottom left) Motion was defined as the change in luminance. (top right) The auditory envelope was averaged within each TR based on the broadcast volume. (bottom right) These variables, along with vocal pitch, were included as sensory regressors in analyses related to pupil area changes. c) Pupil area (averaged for each TR) increased after possession changes, with the solid line and error bars indicating across-subject mean ± SEM and dashed lines indicating each subject. Pupil area change, computed by subtracting a post- (0 to 2 s) minus pre-possession (−6 to −2 s) time period, significantly increased across the boundary. d) Variables entering a mixed-effects model are shown for one subject. We found that surprise significantly predicted pupil area change. e) Separate models run at each time point revealed that surprise predicted pupil area from 0-5 s. Error bars indicate the standard error in the beta estimate. †: 0.05 < FDR-corrected *q* < 0.10. *: FDR-corrected *q* < 0.05. **: FDR-corrected *q* < 0.01.

As expected, subjects’ pupils dilated following possession changes (*p* < 0.001; Fig 4C). More importantly, if pupil dilation reflects event segmentation, pupil dilation across an event boundary should scale with surprise at that boundary. To address this, we asked how surprise, along with the above regressors, predicted pupil area change across possession boundaries from all games using a linear mixed-effects model with subject as a random effect. In line with EST, surprise significantly predicted pupil dilation (*p* < 0.001; Fig 4D; see Table S2 for full details). Additionally, both luminance metrics negatively and the auditory envelope positively predicted dilation. Pupil area change also increased with less time remaining (towards the end of the game). Given the significance of surprise in this analysis, we also ran a follow-up model looking separately at belief-consistent and belief-inconsistent surprise with all of the other regressors (except surprise); we did not have an *a priori* hypothesis about how these subtypes of surprise would relate to pupil dilation. We found that both belief-consistent and belief-inconsistent surprise significantly predicted pupil area change (*p* < .0001 Bonferroni-corrected, in both cases; see Table S2 for uncorrected values). We also re-ran mixed effect models for surprise against pupil area (rather than the pupil area change) at each time point (Krishnamurthy et al., 2017). We found that surprise predicted pupil area between 0-5 s after the possession change (FDR-corrected *q* values < 0.05; Fig 4E; see Table S3 for all time points). Taken together, these results suggest that pupil dilation modulations reflected surprise across possessions.

### Surprise correlates with enjoyment and neural activity in reward-related regions

We next examined if and how surprise relates to subjective enjoyment and activity in neural regions associated with reward. Casual fans typically prefer games with high uncertainty and surprise to combat boredom (Gan et al., 1997; Peterson and Raney, 2008). Indeed, in many circumstances, subjects find tasks to be less boring when events are not perfectly predictable (Geana et al., 2015; Wilson et al., 2019). Accordingly, subjects’ enjoyment ratings given after each game correlated significantly with that game’s mean level of surprise (*r* = 0.82, *p* = 0.007; Fig S3), as well as the standard deviation of surprise values in that game (*r* = 0.92, *p* < 0.001).

We also asked whether subjects valued surprise in the absence of direct experience – that is, would they be more excited to watch games expected to contain more surprise? At the end of the experiment, subjects rated their excitement to watch games from the following year’s tournament (2013), starting from five minutes remaining. We computed future expected surprise for each of these games by finding games from the 2012 corpus that had similar win probabilities at five minutes remaining and then summed the total amount of surprise remaining in those games. All subjects preferred watching games with higher expected surprise in the future (mean *r* = 0.61, range: 0.10 to 0.93, *p* < 0.001; Fig S3). Thus, viewers appear to value both the experience and the expectation of surprise.

In addition to considering enjoyment across a full game, we also investigated the neural effects of surprise on a shorter time scale. Reward signals are intimately linked with the activity of dopamine neurons in regions of the brainstem such as the ventral tegmental area (VTA), as well as targets of those neurons, particularly the nucleus accumbens (NAcc). Classically, the VTA (D’Ardenne et al., 2008; Schultz et al., 1997) and NAcc (Cikara et al., 2011; Gold et al., 2019; Rutledge et al., 2010) respond strongly when rewards are larger or earlier than expected (i.e., reward prediction errors, or RPEs). However, the VTA can respond more broadly to variables other than reward (Engelhard et al., 2019), including sensory PEs (Howard and Kahnt, 2018; Iglesias et al., 2013; Takahashi et al., 2017), unexpected events (Barto et al., 2013; Kafkas and Montaldi, 2015), aversive PEs (Matsumoto et al., 2007), changes in hidden belief states (Nour et al., 2018; Starkweather et al., 2018), reward expectation (Kim et al., 2016; Totah et al., 2013), advance information (Bromberg-Martin and Hikosaka, 2011), and stimulus-stimulus learning (Sharpe et al., 2017), all in the absence of (or controlling for) reward.

Subjects reported having a team preference in approximately half of the games (48 ± 6%; Fig S3). Therefore, in addition to looking at unsigned surprise in all games, for games with a stated team preference we could ask whether neural activity was modulated by the valence of the outcome, or “signed” surprise. We characterized NAcc and VTA activity (Fig 5A) using general linear models (GLMs) with regressors for surprise and signed surprise (Fig 5B). These time courses involve mostly zeros with occasional punctate surprise events, meaning that a physiological process that responds uniformly to any surprise or signed surprise event (i.e., not in a graded fashion) could have a positive relationship. Therefore, we also created “binarized” versions of the surprise and signed surprise time courses to capture processes that respond to surprise but do not track the magnitude of surprise (Leong et al., 2017) (Fig S4). Based on the findings reviewed above, we predicted that the NAcc would respond to RPEs (signed surprise), whereas the VTA could respond to RPEs and/or more broadly to unsigned surprise. In NAcc, activity showed a marginally significant correlation with signed surprise (*p* = 0.056, *t*-test of GLM betas against zero) but not unsigned surprise (*p* = 0.61), providing trend-level support for the standard RPE model (Fig 5C) (see Table S4 for full results). By contrast, a more complex pattern emerged from VTA, which responded significantly to unsigned surprise (*p* = 0.002) and to signed surprise (*p* = 0.009). To follow up on the significant finding between surprise and VTA activity, we ran another model replacing surprise with belief-consistent and belief-inconsistent surprise; we did not have an *a priori* hypothesis about how these subtypes of surprise would relate to VTA activity. We found that belief-consistent surprise (*p* = 0.01, Bonferroni-corrected) significantly predicted VTA activity, and belief-inconsistent surprise showed a trend towards predicting VTA activity (*p* = 0.08, Bonferroni-corrected; see Table S4 for uncorrected values). In sum, we obtained trend-level support for the classic finding that NAcc responses correlate with RPEs using a novel, passively-viewed naturalistic stimulus, whereas VTA responses showed both a classic RPE response and also correlated with unsigned naturalistic surprise.

**Fig 5:**
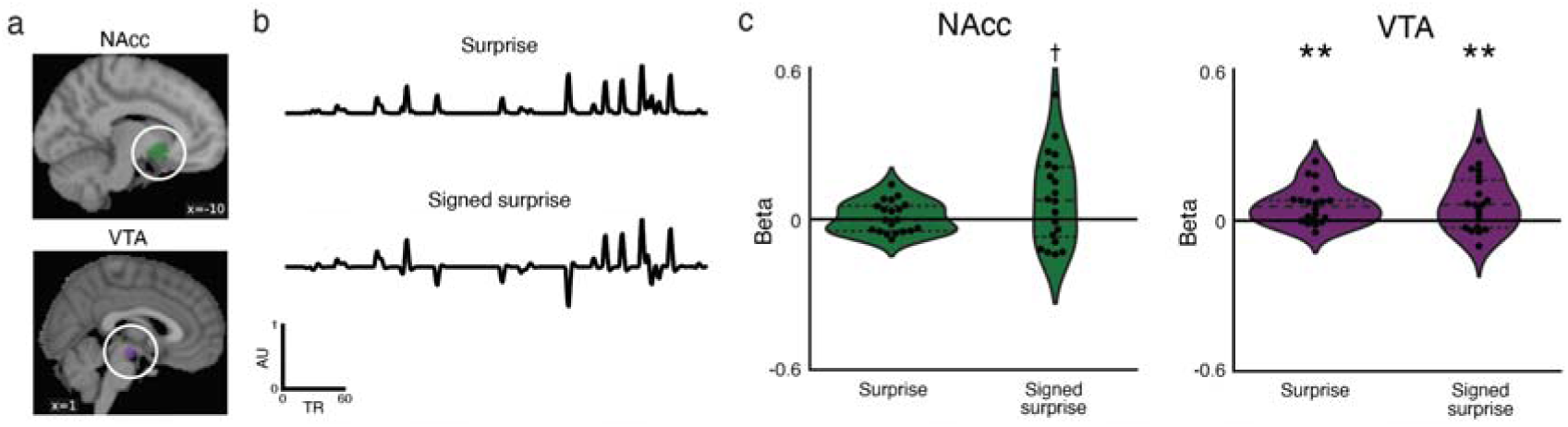
Surprise correlates with neural activity in reward-related regions. a) NAcc and VTA ROIs. b) Depictions for one game of different models included in general linear model analyses (GLMs) for how surprise relates to neural activity in reward-related regions. For all games, we modeled (unsigned) surprise and signed surprise (signed for games with a preference). c) The NAcc responded only in a signed fashion to positive events (top). The VTA responded positively to both unsigned surprise and signed surprise. †: *p* = 0.05. **: *p* < 0.01.

### Surprise and pupil area change positively predict memory for possessions

Numerous studies have found that the segmentation of ongoing experience into discrete events plays a powerful role in organizing memories (Clewett et al., 2019b; DuBrow and Davachi, 2013, 2014, 2016; Ezzyat and Davachi, 2011) and enhances memory for information near the event boundaries (Clewett et al., 2019b; Newtson and Engquist, 1976; Rouhani et al., 2020; Swallow et al., 2010). Given that surprise helps create event boundaries (Franklin et al., 2020) and enhances memory in laboratory settings (Jang et al., 2019; Pine et al., 2018; Rouhani et al., 2018, 2020), we asked how surprise and other factors described above predict long-term memory in our naturalistic paradigm.

We assessed memory by computing the number of possessions subjects recalled with enough specific details that they could be readily identified by an independent rater (Fig 6A). Overall, subjects recalled few possessions (12.0 ± 2.8 out of 157), likely due to high interference given the similar content in each clip. Additionally, subjects’ recall showed temporal contiguity with a powerful forward asymmetry (Howard and Kahana, 2002); all of the recalls transitioned in the forward direction (i.e., if subjects just recalled a possession, the next possession they recalled was from later in the game), and more than 50 percent of the total transitions were to the very next possession (Fig 6B; see Fig S5 for a sample subject).

**Fig 6:**
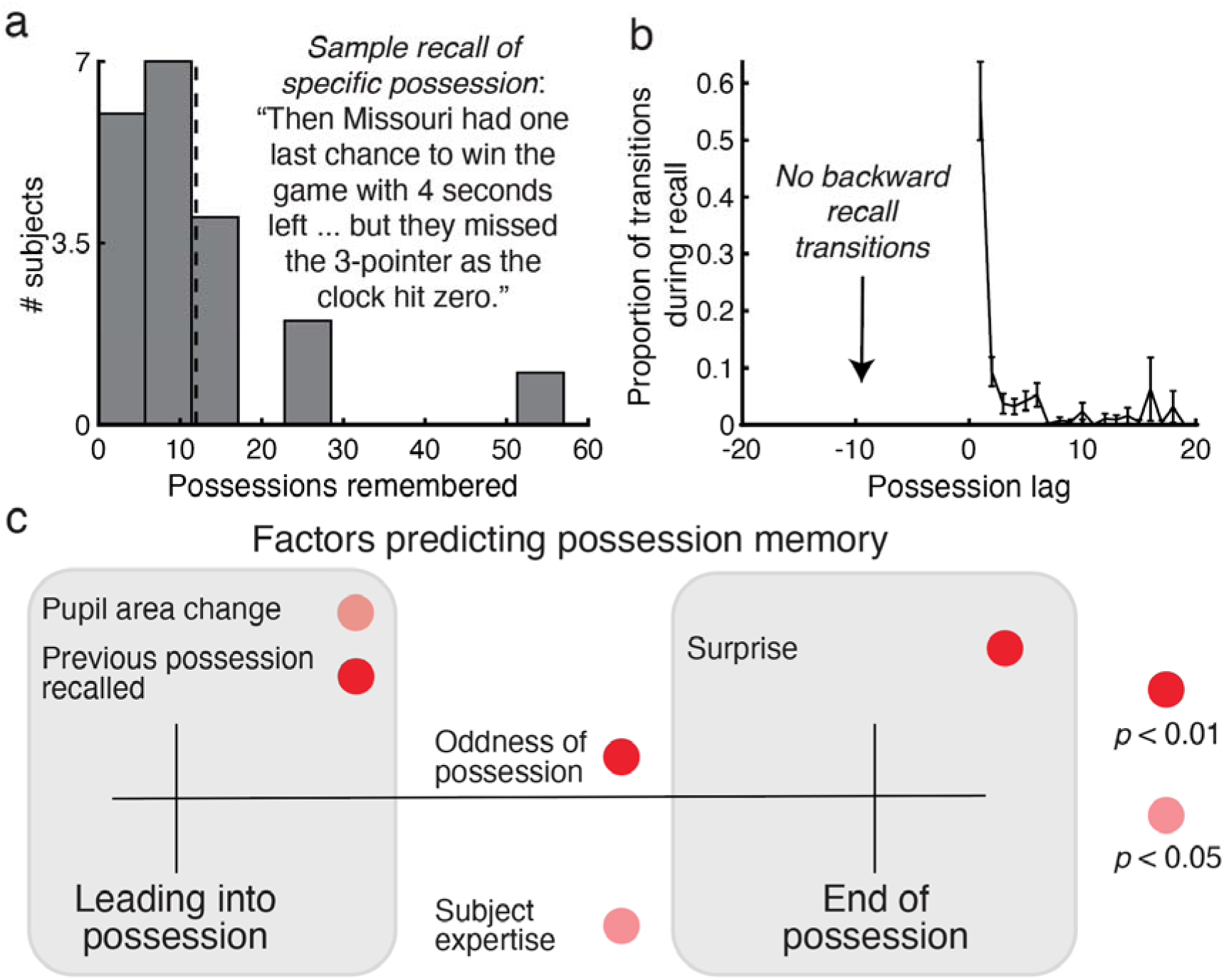
Effects of surprise and physiological and neural factors on memory. a) Sample recall of a specific possession, and the distribution (across subjects) of the number of possessions that were remembered. Vertical dashed line indicates the mean. b) Transitions between successively recalled possession memories showed that subjects overwhelmingly recalled possessions in a forward manner from possession to possession. Error bars represent SEM. c) Schematic of factors that predicted memory in a mixed-effects logistic regression model; a wide range of variables predicted memory for possessions, including pupil area increases leading into a possession, subject expertise, (independently rated) oddness of a possession, and surprise at the end of a possession. Note that all of these significant relationships were in the positive direction.

Theoretically, surprise and other factors occurring at possession boundaries could affect memory for the upcoming possession, or memory for the possession just completed – that is, proactive or retroactive enhancement, respectively. Our first prediction was that surprise at the end of the possession would influence memory. This prediction is based on prior studies that found a relationship between neural activity at the end of an event and subsequent memory for the just-completed event (Baldassano et al., 2017; Ben-Yakov et al., 2013). Indeed, when we took the memorability of each of the 157 possessions (i.e., the average number of participants recalling that possession) and correlated it with end-of-possession surprise, the correlation was significant (*r* = 0.34, *p* < 0.001, permutation test circularly shifting possessions within each game).

The VTA has been linked to storage of long-term episodic memories (Adcock et al., 2006) and peak VTA activation could predict memory on a trial-by-trial basis for the just-ending possession (see Fig S6). However, surprise and/or other factors leading into the possession could also influence memory for subsequent events. For instance, surprise re-orients attention (Itti and Baldi, 2009) and increases subsequent learning rates (Courville et al., 2006; Pearce and Hall, 1980); a recent study also found that prediction error shifts hippocampal connectivity in a manner that should up-regulate encoding (Bein et al., 2020). Pupil dilations also predict higher learning rates (Browning et al., 2015; Nassar et al., 2012; Silvetti et al., 2013). As such, many factors could influence long-term memory of events in the game.

We thus submitted the above factors to a mixed-effects logistic regression model predicting possession memory. We looked at memory for each possession, including separate versions of these factors for the boundary going into the current possession and the boundary at the end of the possession (e.g., we used both surprise at the start of a possession and surprise at the end of the possession as separate predictors of memory for that possession). Significant results are shown schematically in Fig 6C and the full results are detailed in Table S5.

Critically, pupil area changes across the prior possession boundary predicted subsequent memory for the upcoming possession, in line with attentional reorienting and/or an increased learning rate. VTA activity did not predict memory. Surprise at the end, but not the beginning, of the possession predicted memory for the possession. Additionally, we included the following factors, all of which predicted memory individually and in the full model: subject expertise, based on the number of games watched across one’s lifetime, highlighting the importance of domain expertise for memory (Chi, 1978); the “oddness” of a possession, based on an independently-rated assessment of unusual basketball plays (e.g., a lane violation during a critical free throw), which could index “surprise” in a manner not captured by our simplified win probability metric; and, inspired by the temporal contiguity effects depicted in Fig 6B, whether or not the previous possession was recalled. Given that surprise at the end of the possession predicted memory, we also ran a follow-up model removing surprise at the end of the possession and including belief-consistent and belief-inconsistent surprise; we did not have an *a priori* hypothesis about how these subtypes of surprise would relate to memory. We found that both significantly predicted memory (*p* < .01, Bonferroni-corrected; see Table S5 for uncorrected values). Altogether, these results capture complex, multifaceted aspects of real-world memory and highlight the importance of surprise, among other associated factors, in shaping those memories.

## Discussion

Our findings reveal that the popular activity of competitive sports viewing is a valuable example of naturalistic surprise, and our analyses of this task led to multiple novel behavioral and physiological discoveries in support of the tenets of event segmentation theory (EST) (Franklin et al., 2020; Zacks et al., 2011). Namely, surprises appear to strongly coincide with the segmentation of internal event representations, indexed by increased subjective perception of event boundaries, increased pupil dilation, an increased likelihood of significant neural representational shifts (as measured using a Hidden Markov Model), and increased subsequent memory for events.

Further results provide novel evidence for reinforcement learning models of reward prediction errors (RPEs) in a naturalistic, passive-viewing setting. In the nucleus accumbens, we observed a trend towards classic RPE (signed surprise) effects, reflecting dynamic changes in the probability that a preferred team would win a basketball game (Cikara et al., 2011). In the ventral tegmental area, we also found classic RPE effects, as well as activity correlated with unsigned updating of beliefs, extending previous work on the VTA to a naturalistic setting (Howard and Kahnt, 2018; Sharpe et al., 2017; Starkweather et al., 2017, 2018; Takahashi et al., 2017). Finally, in showing that surprise correlates with subjective enjoyment, we provide further support for the intriguing idea that – when information is not instrumental for survival – humans may actually prefer unpredictable scenarios (Ely et al., 2015; Geana et al., 2015).

Importantly, our approach illustrates a dissociation of surprise effects based on whether they bolstered or contradicted the currently held predominant belief in who would win. Belief-inconsistent surprise – which also leads to a game state of higher uncertainty (Nassar et al., 2019; Shin and Dubrow, 2020) – was a significantly better predictor of subjective event boundaries than belief-consistent surprise. Furthermore, transitions between neural states in precuneus and mPFC were significantly correlated with belief-inconsistent surprise but not belief-consistent surprise, and the correlation with belief-inconsistent surprise was larger than the correlation with belief-consistent surprise for both regions. Contrary to the above results showing a distinction between belief-inconsistent and belief-consistent surprise, measures such as pupil dilation, activity in VTA, and memory showed significant or trending effects for both belief-consistent surprise and belief-inconsistent surprise. Ultimately, the fact that different flavors of surprise have different behavioral, physiological, and neural outcomes demonstrates that individuals’ predictions may have both a binary aspect (i.e., which team will win?) and a probabilistic one (i.e., how likely is it?) (Johnson et al., 2020). These discrepancies raise questions about how and where these two putative aspects of predictions diverge, opening up avenues for future research.

Here we have defined surprise as a change in the estimated outcome of the game. This definition appears on its surface to conflict with definitions used in a prior report (O’Reilly et al., 2013), which designated events that signal updates to an underlying probability distribution as “model updates” and errors in predicting moment-to-moment events as “surprises”. We agree that surprise can reflect expectation violations stemming from momentary events. In a basketball game, that could take the form of a player making a truly spectacular and improbable shot, which might incidentally have the same influence on win probability (and therefore, our measure of surprise) as a regular shot. However, we argue that higher-level updates about the world (e.g., who will win the game) can also be surprising. For instance, making a 3-point shot that increases a team’s likelihood of winning by 79% is far more surprising (in the common-use sense of the term) than an identical play that only increases the likelihood by 12%, even though the likelihood of the event itself (e.g., making the 3-point shot) is identical; most sports fans would characterize these kinds of major shifts in win-probability as surprising, and they are typically accompanied by strong emotion (e.g., shock, elation, disappointment). We use “surprise” here to underscore the point that updating an internal belief state by a large amount at a single time point likely shares many subjective features with witnessing an improbable momentary event; in both cases, there is a discrepancy between what one previously believed would happen (that team A would win the game; that the shot would not go into the basket) and what follows (now you think team B will win; you see the ball go in the basket).

Another key conceptual issue is whether “surprise”, defined as above, should be viewed as a construct in the world (i.e., a change in the likelihood of one team winning, according to the mathematical model devised by the experimenters) or a construct in the subject’s mind (i.e., a change in the subject’s belief about which team will win). While these are clearly distinct concepts, we believe that they coincide in our study. That is, our approach is based on the hypothesis that our (basketball-savvy) subjects have a predictive model in their head that corresponds to the model that we (the experimenters) are using to predict the outcome of the game – as such, moments when our regression model shows a change in the win probability correspond to moments when subjects show a change in their (internal) win probability model. Of course, this hypothesis could be incorrect -- but the strong correspondence between the predictions of our win-probability model and observed neural, behavioral, and physiological data lends converging support to this hypothesis.

A related conceptual issue is the relationship between “surprise”, defined as above, and event segmentation. Event Segmentation Theory (Zacks et al., 2011) and related perspectives (Clewett et al., 2019b; Franklin et al., 2020; Shin and Dubrow, 2020) posit a cascade of cognitive processes that occur when you make observations that are incompatible with your current understanding of the world (e.g., flushing of your current representation of the situation, loading up of a new and more suitable predictive model). In these models, the process of event segmentation starts when your current understanding of the world is destabilized by a new observation and ends when you settle on a new understanding. Viewed from this perspective, high levels of surprise (defined here as substantial changes in the subject’s belief about the game outcome) and event segmentation are two sides of the same coin. When we say that an event is “highly surprising”, this refers to the substantial change in belief triggered by the new observation, and “event segmentation” refers to the underlying cognitive machinery that supports this change in belief. In our study, we operationalized event segmentation by having subjects press a button when they thought one meaningful unit ended and the other began (Newtson, 1973). We believe that subjects followed these instructions by noticing when their internal belief state changed and then pressing the button – so, by this logic, noticing the surprise (the change in beliefs) prompted subjects to make a button press.

Lastly, one noteworthy aspect of our investigation is that the probabilistic predictions derived from our model can be validated behaviorally (as shown in our prediction test, Fig 1E). We speculate that similar latent belief states underlie people’s responses to real-world events in other domains, including fiction, film, and the news – people are elated (or dismayed) in proportion to their surprise at breaking news stories or sudden narrative swings and are more likely to consume exciting forms of media with many twists and turns. Moreover, people’s most long-lasting memories are formed in precisely those moments when their beliefs substantially shift. Future studies should continue to leverage naturalistic stimuli with quantifiable latent variables to investigate how humans respond to their ever-changing world.

## Supporting information

All supplemental figures.

## Acknowledgments

This work was supported by a CV Starr fellowship to JWA and the ONR MURI grant N00014-17-1-2961 to KAN and UH. We thank Wazee Digital and the NCAA for game footage, Vishnu Murty for help with ROIs, Lisa Musz for the free recall scoring rubric, James Howard and Jeff Zacks for helpful comments on drafts of this manuscript, and Chris Baldassano, Kelly Bennion, Silvy Collin, Nick Depinto, Robert Hawkins, Manoj Kumar, Qihong Lu, Rolando Masis-Obando, Lizzie McDevitt, Anne Mennen, Sebastian Michelmann, Ida Momennejad, Sam Nastase, Mark Pinsk, Victoria Ritvo, Nina Rouhani, Monika Schönauer, and Jamal Williams for assisting with data collection and/or various aspects of this project.

## Author contributions

J.W.A. conceived the experiment. S.D.M., U.H., and K.A.N. contributed to the experimental design early on and provided numerous analysis ideas. J.W.A. programmed the experiment, collected some of the data, and performed the bulk of the analyses. T.H. scored the recall data. K.P. computed and provided all win probability metrics. J.W.A., S.D.M., & K.A.N. wrote the manuscript. All authors discussed the results and revised the paper.

## STAR Methods

### RESOURCE AVAILABILITY

#### Lead Contact

Further information and requests for resources and reagents should be directed to and will be fulfilled by the Lead Contact, James Antony.

#### Materials Availability

The basketball videos have restrictions due to copyright reasons. However, much of the metadata, including everything used for the analyses herein, are included with the openly available data (see “InGameVars.xls”). Please e-mail the Lead Contact for other information about the materials.

#### Data and Code Availability

All code and data from this project will be made openly available upon publication on Princeton’s Dataspace. Neuroimaging data will be shared on OpenNeuro.org in the brain imaging data structure (BIDS) format amenable to meta-analyses and reproducible neuroscience.

### EXPERIMENTAL MODEL AND SUBJECT DETAILS

#### fMRI subjects

Twenty subjects (6 female, 18-35 years old) with normal or corrected-to-normal vision and fluent in English were recruited via campus flyers and word-of-mouth. Subjects were given hourly monetary compensation for participating ($20/hr). Written informed consent was obtained in a manner approved by the Princeton University Institutional Review Board. All subjects self-professed to having seen or played in more than 50 basketball games (across all competitive levels of the sport).

### METHOD DETAILS

#### Stimuli

The last five minutes from all 32 Round-of-64 2012 NCAA® tournament games were acquired as audiovisual files for a fee and with permission from Wazee Digital™. These files were down-sampled to visual dimensions of 1280 × 720 and frame rate of 29.75 Hz and audio dimensions of 48,000 Hz for computational efficiency when presented using the MATLAB Psychtoolbox software. These games were additionally edited to reduce their overall length by eliminating breaks in action (other than brief intervals preceding free throws and in-bound passes) in a manner that did not significantly compromise their overall comprehensibility, resulting in clips between 5:29 and 7:32 in length. Tournament games from the Round-of-64 were used for the following reasons: tournament teams are given “seeds” that inform subjects about the teams’ relative strengths (stronger teams have lower numbered seeds, so #1 seeds are the strongest teams), which should aid subjects’ win probability estimations; tournament games have a heightened sense of importance relative to the regular season, enhancing subject engagement; the Round-of-64 is an early round of the tournament that we intuited subjects would be unlikely to remember (if they had seen or read about the original broadcasts). Nine games were selected for presentation in the scanner using the following criteria: Games were selected to have as wide-ranging amounts of surprise as possible (Fig S1); games were also selected to have as wide-ranging tournament seeds as possible, except that no games between #1 and #16 seeds were selected, as no #1 seed had ever lost before 2012 – each of the other types of matchups (#2 versus #15, #3 versus #14, #4 versus #13, #5 versus #12, #6 versus #11, #7 versus #10, and #8 versus #9 seeds) was selected at least once; games involving extremely well-known teams (e.g., North Carolina) were given a lower priority to reduce the likelihood that a subject may have remembered the outcome. The selected games/seeds were as follows: #2 Missouri versus #15 Norfolk State; #3 Florida State versus #14 Saint Bonaventure; #4 Indiana versus #13 New Mexico State; #5 Wichita State versus #12 Virginia Commonwealth University; #6 University of Nevada-Las Vegas versus #11 Colorado; #7 Gonzaga versus #10 West Virginia; #7 Saint Mary’s versus #10 Purdue; #7 Notre Dame versus #10 Xavier; #8 Creighton versus #9 Alabama.

#### Design and procedure

The experiment included four phases: game viewing, free recall, prediction test, and anticipated surprise preference test (Fig 1). Subjects arrived in the scanner suite, learned about MRI safety, gave informed consent to participate, listened to the instructions, and then entered the magnet.

The main section of the experiment consisted of three alternations between game viewing and free recall phases. Game viewing phases included three games in succession as part of one scanning run. To keep run length approximately even, the nine videos were pseudo-randomly shuffled so that one of the longest three, one of the shortest three, and one of the remaining three were shown in each run. Games were presented in the middle 80% of the screen so the subject’s view would not be obstructed. Audio was delivered via scanner earbuds and volume levels were tested before the games using an internet video. Audio from the regular broadcast, including crowd noise and commentary, was included to maintain a naturalistic viewing experience. Subjects were asked to simply watch the games, though they were told beforehand that they would later be asked to freely recall the games using as much specific detail as possible. After each game, they were asked to indicate the team for which they were cheering or if they had no preference. We emphasized that they should not feel obligated to prefer one team or another. After this question, they indicated how exciting they found the game from 1 (not exciting) to 7 (exciting).

During recall phases, subjects freely recalled the three previously seen games (one at a time, in the order in which the games were viewed) as part of one scanning run. Subjects were shown the names of the two teams involved in the game and were asked to recall the game in as much detail as possible. For instance, they were asked to include the score and approximate amount of time left during any possession they could remember and any parts of that possession as they unfolded (e.g., a screen, a pass, a drive, a defender’s move, the outcome of a shot).

After three rounds of viewing and free recall, subjects remained in the scanner to receive anatomical and field map scans while they performed two more behavioral tasks. In the first, subjects were asked to rate the likelihood of teams winning for a series of possessions from five games (Fig 1E). These games were also taken from the 2012 tournament and were chosen to evenly distribute win likelihoods and resemble the likelihood distributions of the viewed games as much as possible (Fig S1). Subjects were shown the teams, their seeds (i.e., their relative strengths), their scores, which team had the ball, and the amount of time remaining, and they were asked to slide a joystick from 0-100% to indicate the likelihood that the stronger-seeded team would win. These scenarios were updated after every possession, so each game had multiple ratings. The selected games/seeds were as follows: #2 Duke versus #15 Lehigh; #4 Michigan versus #13 Ohio; #5 New Mexico versus #12 Long Beach State; #6 Cincinnati versus #11 Texas; #8 Memphis versus #9 Saint Louis. We correlated these likelihoods against the algorithm to assess prediction abilities. Data were omitted from the few instances in which subjects knew the outcome.

In the second task, subjects rated (from 1-7) how excited they would be to watch 28 games starting with five minutes remaining from the Round-of-64 in the 2013 tournament (Fig S3B). All games were selected, except for games between #1 and #16 seeds for the reasons delineated above. We found the anticipated level of surprise by finding games from the 2012 regular season corpus with similar win probabilities with five minutes remaining and calculating the actual amount of surprise remaining in those games. Then, we correlated these excitement ratings with anticipated surprise for each subject (Fig S3B).

After these tasks, subjects left the scanner and completed a questionnaire. Questions included approximately how much basketball they had watched (i.e. their expertise), their most and least favorite teams (none were involved in the games they viewed except for one subject’s third favorite team), whether they knew or suspected the outcomes of any games that they viewed or that were part of the prediction tests (two subjects knew the outcomes for one viewed game each; two subjects knew the outcome of one game in the belief test), their level of overall engagement (6.55±0.25 out of 10), and whether they found any parts of the games particularly surprising (two subjects mentioned an unusual lane violation during a critical free throw).

#### Win probability metric

To calculate win probabilities for each possession, we created a model based on four factors: the difference in score between the teams oriented to be positive when the “home” team is winning and negative when they are losing, the amount of time remaining, which team is in possession of the ball, and, where publicly available, an adjustment based on team strength. This model was trained using data from every regular season game from the 2012 season. Using the first half of the corpus data, we found every instance of a particular “game state”, binned by every score difference from −20 to +20 for home - visitor score, every time bin within 6 seconds (e.g., 60-54 seconds left in the game), and either team possession (home or visiting team). From these instances, we “peek ahead” and compute the percentage of instances in which the home team won that game to create a home win probability for that bin. Following the tendency of a basketball analyst’s algorithm (see below) to avoid making predictions of absolute certainty, we added up to 0.5% of noise for any state with exactly 0% or 100% probability (probability values of 0% were replaced with a value uniformly sampled between 0.0% to 0.5%, and probability values of 100% were replaced with a value uniformly sampled between 99.5% to 100.0%) Then we applied light smoothing across bins, with no smoothing for the final two time bins to avoid having too much blurring in scenarios with extremely well-defined probabilities. In Fig 1B, we plotted time bins on the x-axis and score differences on the y-axis with win probability in color from 0-100. We created separate plots for when the home team possesses the ball (top), when the visiting team possesses the ball (middle), and – because it can be difficult to spot the differences in the top two plots – we created a third plot showing the difference between them, i.e., the difference between the home team having possession and the visiting team having possession, controlling for the same time / score difference bins (bottom).

In observing the model, first and foremost, score difference strongly predicts win probability. Second, this advantage relies on time, as the score difference has a larger effect on win probability as time elapses. As an example, having a 5-point lead with 300 seconds left (∼70% win probability) is less advantageous than the same lead with 5 seconds left (∼99% win probability). Third, the advantage for having possession of the ball is relatively small for much of the game, but becomes quite large in a close game with very little time left.

We validated the model’s win probabilities by training on the first half of corpus games and testing on the second half in two ways: 1) against win probability estimates from an expert sports analyst and 2) against the actual outcomes from an out-of-sample subset of games. We obtained win probability estimates created for a basketball analytics website using a custom, proprietary algorithm by Ken Pomeroy (K.P.; www.kenpom.com). These estimates were created using similar factors, such as the score difference between the teams, the amount of time left, the team in possession of the ball, and the strength of the two teams. We obtained these data from the entire 2012 season via personal correspondence, although the data are also available on K.P.’s website. Our estimates on the second half of games show a strong correspondence with those of K.P. (r = 0.998) (Fig 1C, top). We also compared the predictions of our model to the true likelihood on the second half of the data. The 4-factor model also showed a strong correspondence with these likelihoods (r = 0.9992).

For the final version of the model to use for the viewed games, we trained the model again using *all* of the regular season games. Then we added the final factor to the model: an adjustment based on the relative strength of the teams obtained from K.P.’s publicly available website as the score difference in the predicted outcome. For example, Missouri was expected to beat Norfolk St. by 23 points (86-63). From this, we can compute how much we expect Missouri to outscore Norfolk St. on a per second basis [23 points / (40 minutes in a full game * 60 seconds / min) = + 0.0096 points per second]. We then created an “expected score difference” value for each possession by adding [e.g., 0.0096 * the number of seconds remaining in the game] to the actual score difference and used this metric in the model to find the win probability. We validated this final model against K.P.’s predictions on the viewed games (Fig 1C, bottom) (*r* = 0.996).

#### Surprise calculation and related metrics

As an agent traverses the world, they proceed through a series of states. In viewing basketball, one could conceptualize a momentary state as the current score, the team in possession of the ball, the relative strength of the two teams, and the amount of time left in the game. In each state, an ideal observer, via repeated experience (e.g., watching games), could form a refined, probabilistic belief in some outcome (e.g., which team will win) via an iterative process like temporal difference learning (Sutton and Barto, 1998).

As formulated elsewhere (Ely et al., 2015), surprise is the change in prediction from the previous to the current state (here, the change in win likelihood). For the purposes of aligning surprise as a psychological phenomenon with physiological and neuroimaging data, surprise was labeled as ‘0’ for all stable time points (e.g., within a possession) and the magnitude of the surprise scaled with changes in belief state (resembling a “stick” function with spikes of varying magnitudes; Fig 1D).

Some analyses separated “belief-consistent” from “belief-inconsistent” surprise. To do this, we first found the team with a higher than 50% chance of winning. Events that increased this probability were classified as belief-consistent, and events that decreased it were classified as belief-inconsistent. There were no differences between the mean size of surprise for these constructs (belief-consistent: 6.44 ± 0.12; belief-inconsistent: 6.54 ± 0.08, *p* = 0.93).

#### Free recall scoring

Subject recall recordings were sampled at 11,025 Hz using built-in MATLAB software and converted for transcription. Each TR in which subjects spoke was classified as belonging to one of the following recall types: (1) veridical recall, with sufficient specific detail to be identified as a particular possession, (2) gist-based, summarized recall that is temporarily imprecise but accurate, (3) inaccurate detail that is nonetheless about the game, (4) irrelevant commentary about the game, and (5) words that are not related to recall. Additionally, for each TR in which subjects spoke, we recorded: which (if any) game was being recalled; if (1) for recall type, which possession is being recalled, or if (2), which are the first and last of the numerous possessions that are being summarized; if (1) for recall type, does the recall relate to the first or the second half of the possession? After creating these labels, we considered for a given subject only whether each possession was recalled with specific detail (1) and the order of these recalls with respect to the order in which they occurred in the game.

#### Memory metrics

We calculated the temporal contiguity effect (Fig 6B) by finding, for each position after the initial one, the lag (in number of possessions as they actually occurred) with respect to the prior recalled possession. We then normed lags within-subject so that every subject’s proportions summed to one. See Fig S5 for one subject’s possession recall transitions.

For the mixed effects logistic regression analysis predicting memory across possessions, we ran a model including all of the following factors: surprise, pupil area change, and peak ventral tegmental area (VTA) activation for transitions leading into the possession and at the end of the possession, previous possession memory, oddness of possession, and subject expertise. All analyses for factors leading into possessions had no data (NaN values) for the first possession of the video clip. Peak VTA activation was calculated as the value at each possession at its peak of −2 s minus a baseline averaged from −12 to −8 s (Fig S6). “Oddness” of possessions was determined independently by an expert for unusual basketball plays (e.g., a lane violation during a critical free throw). Only 2 out of 157 possessions were labeled this way. Subject expertise was coded as 1-3 depending on the number of games played in/watched over the lifetime (8 subjects had watched 50-200 games; 9 subjects, 200-1000 games; 3 subjects, over 1000 games). Significant factors were depicted in Fig 6C and a full report of the analysis can be found in Table S5.

#### Event segmentation behavioral experiment

15 subjects (8 female, 18-35 years old) with similar characteristics were recruited as above and were given hourly monetary compensation for participating ($10/hr for behavioral studies). Written informed consent was similarly obtained. Subjects watched the same games as those viewed in the scanner, though in this case they watched all 9 games consecutively. They performed the following task as they watched: “Click the mouse when, in your judgment, one unit of the game ends. Mark off the game you’ll be seeing into the largest units that seem natural and meaningful to you.” (Newtson, 1973) These subjects performed the prediction test, but did not perform recall nor the future surprise preference test. We labeled mouse clicks occurring within 2 seconds after a possession change as an endorsement of it as an event boundary and aligned these responses with other possession-level metrics (Fig 2A). Specifically, for each boundary, we computed a *subjective boundary agreement* score, quantifying the agreement across subjects (ranging from 0 for no agreement to 1 for perfect agreement) that a game unit ended at that possession boundary.

#### Physiological measurements and analysis

Eye tracking data were acquired in the scanner at 1000 Hz using EyeLink 1000 software (SR Research, Inc., Mississauga, Ontario Canada). Eye tracking data from 6 subjects were lost due to technical difficulties. Extreme outliers (z > 5) and time periods when the gaze location was off-screen were removed, and after exclusion missing pupil area and eye location data were interpolated by averaging around the errant time periods.

Pupil area measurements are influenced by both gaze location and video luminance. Normally, experimenters measuring gaze location only present stimuli centrally or in fixed locations (e.g., Eldar et al., 2013); however, we reasoned that restricting subjects’ gaze while watching videos would artificially affect their viewing experience. Instead, we accounted for gaze location confounds by binning x- and y-coordinates and z-scoring pupil area within each bin (Fig 2A). We addressed confounds related to video luminance by calculating the mean global gray-scaled luminance for the entire screen and local gray-scaled luminance corresponding to the approximate visual angle of the fovea (2°) (Choplin and Edwards, 1998) surrounding the current eye location for each second of video. We addressed confounds related to video motion by calculating the frame-by-frame change in luminance for every pixel, creating a difference image (Puttegowda and Padma, 2016). For global motion, we averaged these pixel-based changes across the full screen, and for local motion, we averaged these changes within 2° of the current eye location. We addressed the confound of auditory volume by averaging the auditory envelope of the broadcast for each second of video and the confound of commentator prosody by finding the fundamental frequency (f0) in the audio stream using Praat software (http://www.fon.hum.uva.nl/praat/) (Boersma, 2001). Brief stretches without speech were given NaN values and were omitted from the regression. We addressed possible confounds relating to the position of the ball on the court by creating an experimenter-annotated regressor indicating whether the ball was on the left or right side of the game telecast. We also added a regressor indicating the amount of game time remaining at the start of each possession. Regressors were averaged and down-sampled to one value per second.

For time-course analyses locked to possession boundaries, normed pupil area was averaged within each second and then across trials. A *t*-test was then performed across subjects between the pre (−6 to −2 s) and post interval (0 to 2 s). To relate pupil area measurements across possession boundaries to surprise, we entered the amount of game remaining, global luminance, local luminance, global video motion, local video motion, the auditory envelope, commentator prosody (f0), court position, and surprise into a mixed-effects, linear regression model predicting pupil area change using subject as a random effect. Global luminance, local luminance, global video motion, local video motion, the auditory envelope, and commentator prosody (f0) were entered as their change values from average post – average pre values, whereas court position and game remaining were entered as their value at the beginning of the new possession (since the subtraction would be theoretically meaningless). We also re-ran separate linear, mixed-effect models looking at how surprise across the possession boundary predicted pupil area (rather than pupil area change) at every time point in the interval. For this analysis, we applied FDR correction assuming dependence across tests because of the temporally autocorrelated pupil signal (Benjamini and Yekutieli, 2001).

#### FMRI acquisition and preprocessing

Neuroimaging data were acquired on a 3T full-body Siemens Prisma scanner with a 64-channel head coil, using a T2*-weighted echo planar imaging (EPI) pulse sequence (simultaneous multislice factor 4, no in-plane acceleration, TR 1000 ms, flip angle 59°, TE 30 ms, whole-brain coverage 56 slices of 2.5 mm thickness, in-plane isotropic resolution of 2.5 mm, FOV 195 mm). The first preprocessing steps were performed using FMRIprep (https://github.com/poldracklab/fmriprep; Esteban et al., 2019), including motion correction, susceptibility distortion correction (using field maps or the “use-syn-sdc” flag in their absence), brain tissue segmentation, and coregistration and affine transformation of the functional volumes to the 1 mm isotropic T1w anatomical and subsequently to MNI space.

The data were imported, down-sampled to a 3 mm isotropic resolution and three scans from the beginning of each run were discarded. Next, the data were smoothed using SUSAN smoothing (https://fsl.fmrib.ox.ac.uk/fsl/fslwiki/SUSAN, Python code adapted from: https://github.com/INCF/BrainImagingPipelines/blob/master/bips/workflows/gablab/wips/scripts/modular_nodes.py) with a 5-mm full width-half maximum spatial kernel. Next, the data were masked using an across-run averaged mask, z-scored, high-pass filtered (140 s cutoff) (Aly et al., 2017; Chen et al., 2017), and confound variables [movement in three directions, rotation in three directions, framewise displacement, and six anatomical components used to correct for the influence of physiological noise (“a_comp_cor_00” to “_05” in fMRIprep) (Behzadi et al., 2007)] were regressed out at the run level. Data from the first 10 s of each video were removed to avoid confounds related to strong onset responses (Nastase et al., 2019).

#### Regions of interest (ROIs)

Binarized V1, precuneus, and mPFC ROIs were obtained from a previous dataset (Simony et al., 2016). V1 indicates early visual cortex and nearby voxels; the ROI was created using voxels near the calcarine sulcus with strongest inter-subject correlation. Precuneus and mPFC were calculated using whole-brain functional connectivity using posterior cingulate cortex as the seed, separating the default mode network into 10 parts with significant correlations and labeling the masks based on cluster location. The NAcc ROI was obtained using an association test in Neurosynth (threshold: *z*>10) based on the term, “nucleus accumbens”. A probabilistic ventral tegmental area (VTA) ROI was obtained from (Murty et al., 2014) via personal correspondence, and a 75% probability threshold was used to binarize it, resulting in 37 voxels. Additionally, HMM analyses were performed using parcels from the Harvard-Oxford Brain Atlas package from FSL 5.0 (Smith et al., 2004).

#### Hidden Markov model (HMM) analyses

The start of each video was locked to a pulse from the scanner to align the hemodynamic response for each moment in the video across subjects. In an additional preprocessing step, the imaging data were shifted 5 s (5 TRs) for later alignment with marked events.

All HMM analyses were performed using the Brainiak toolbox function, brainiak.eventseg.event.EventSegment (Kumar et al., 2020). Each HMM state was composed of a particular mean activity pattern across all voxels within the region, and each instance of a neural pattern was assumed to be normally distributed around this mean. Following prior work (Baldassano et al., 2017), this particular analysis function imposes the constraint that the HMM cannot re-visit a state once it leaves that state. In other words, each new neural pattern is either assigned to the same state as the previous time step, or a new (not-previously-visited) state. HMMs were trained on a version of this Brainiak function that provides better fits when event states are uneven in length (“split_merge=True” in Brainiak v0.10). To train the HMMs within each ROI/parcel, we first found the best-fitting number of states for each game, using a nested cross-validation procedure. On each of the “outer loop” folds for this cross-validation procedure, we selected a single test subject; the other 19 subjects were split into 14 training subjects and 5 validation subjects. Data from the 14 training subjects were averaged together, and data from the 5 validation subjects were averaged together. For the “inner loop” of the cross-validation procedure, we tried versions of the model with different numbers of HMM states ranging from 1 to 30. In each of these “inner loop” folds, we trained the model on the (averaged) data from the 14 training subjects, applied that model to the (average) data from the 5 training subjects, and computed the log-likelihood of the fit to the validation set (for robustness, we used four different ways of dividing the 19 non-test subjects into a training set and a validation set). Based on the “inner loop” results, we chose the number of states that maximized the log-likelihood of the fit to the validation set. We then fit a new HMM to the withheld “test” subject using this number of states. Full details of the basic model are described elsewhere (Baldassano et al., 2017).

We were first interested in where the HMM placed state transitions and how this aligned with possession boundaries and surprise at those boundaries. To address the question of whether the HMM state transitions aligned with possession boundaries, we first averaged the HMM state transition time course (=1 when there is a transition, 0 otherwise) across subjects; we then smoothed both the averaged HMM state transition time course and the possession boundary time course (binary 1/0) by taking running averages using a moving window of ± 7 s; finally, we concatenated the smoothed time courses across all nine games and correlated the smoothed time courses for HMM state transitions and possession boundaries. To assess the strength of these correlations versus chance, we created null distributions in two ways. First, we circularly shifted the possession boundary time course within each game. This means that we shifted the possession boundary time course ahead by a random number of TRs, cut off the part that extended beyond the neural time course, and moved that part to the beginning of the possession boundary time course. This shift was performed 10,000 times to random extents within each game, with the provision that the circular shift could not land within 1 TR of the same time course. Second, we scrambled the order of possessions (preserving possession length) within a game 10,000 times. In both cases, we then compared the true versus scrambled distributions (Fig 3B shows the latter method).

Next, we were interested in whether HMM boundaries increase with surprise, which we computed both across games and possessions. Across games, we correlated the best-fitting number of states per minute for each game with mean surprise in that game, and we compared this with null distributions by shuffling the mean surprise for each game 10,000 times (Fig 3C). Control correlations were also run to verify these correlations could not be explained by the number of possessions per minute or total amount of visual motion per minute (which captures, e.g., camera angle shifts), and linear regression analyses including all of these factors were run to investigate whether the surprise effect survived when these factors were considered. Significance was assessed in a similar manner by comparing the true betas to those in permutation tests.

Across possessions, we asked whether HMM state transitions occurred more often at surprising possession boundaries. To answer this question, we first calculated (for each region of interest, and for each possession boundary) HMM *state transition agreement*, operationalized as the proportion of subjects showing at least one HMM state transition in the 15-s window spanning the possession boundary (±7 s around the boundary) (Fig 3D). There were 157 possessions in total across the 9 games, so we had 157 of these HMM state transition agreement values. Next, we correlated HMM state transition agreement with the amount of surprise at each boundary across the 157 possessions. Finally, we compared this correlation with null distributions whereby we circularly shifted the surprise assigned to each possession within game, with the provision that the new circularly shifted order could not land within 1 of the original possession order. In other analyses, we repeated this procedure for belief-consistent and belief-inconsistent surprise, all against similar null distributions. We also correlated HMM state transition agreement with subjective boundary agreement from the other group of subjects. For HMM analyses applied to parcels from the Harvard-Oxford Brain Atlas, we calculated FDR values for each parcel, assuming independence (Benjamini and Hochberg, 1995).

#### Neural general linear model (GLM) analyses

We created parametric regressors by aligning discrete events with each video. These events were convolved using the canonical HRF to create a time course for the regression. For signed analyses, positive values indicated positive events for the preferred team, and vice versa. When no team was preferred in these analyses, the entire game was modeled as a row of zeros. Data scrubbing (Power et al., 2012) was performed by finding TRs with a framewise displacement above 0.33 and removing the segment starting 1 TR before and 2 TRs after (Clasen et al., 2014). Because our surprise regressors remained at zero in between surprise events, a brain area responding similarly to every win probability change (rather than in a graded, linear fashion) could also result in a positive beta. Therefore, we also created binarized “onset” regressors, whereby all events were given the same binary positive value for unsigned surprise and the same positive or negative value for signed surprise. Under this regime, positive betas for the regular versions will only occur when there is a linear relationship with the amount of surprise rather than just any surprise. The following regressors entered each GLM: the amount of game remaining in seconds of game time at the start of the possession; which team was in possession of the ball; the auditory envelope; global video luminance; global video motion; commentator prosody; court position of the ball (left or right); framewise displacement of the head; (unsigned) surprise; signed surprise; unsigned, binarized surprise; and signed, binarized surprise (Table S4). We also ran a follow-up GLM removing unsigned surprise and including belief-consistent and belief-inconsistent surprise (Table S4). Neural GLMs were performed using the linear_model.LinearRegression function in Python’s scikitlearn toolbox. Note that framewise displacement showed a significant relationship with VTA activity (*t* = 3.83, *p* = 0.001) (Table S4), raising the potential concern that it separately correlated with surprise. We therefore correlated each subject’s framewise displacement on each TR with the time course of surprise. These correlations did not differ from zero, contrasting the Fisher r-to-z transformed correlation coefficient against a row of zeros (*t* = 1.35, *p* = 0.19).

### QUANTIFICATION AND STATISTICAL ANALYSIS

For analyses assessing single-subject correlation coefficients (e.g., the prediction test, the future surprise test), we computed Fisher *r*-to-*z* transformed values before comparing to zeros by paired *t*-test.

For all permutation analyses, we used 10,000 permutations and assessed significance by whether true values fell below the 2.5^th^ or above the 97.5^th^ percentile of the null distribution. For event segmentation and memory analyses, we correlated various surprise measures against across-subject boundary agreement and across-subject possession memory, respectively, and we assessed chance by circularly shifting the possession order within each game while keeping the order of the games intact. Note that this is a highly conservative method for assessing chance, and that null distributions using this metric are occasionally above zero due to across-game differences, because circularly shifted values for a game high in surprise will shift onto other high surprise values more often than chance.

In the HMM boundary time course analyses, we correlated smoothed HMM state transition agreement against a smoothed boundary time course concatenated across all games. We assessed significance by comparing to two types of null distributions: 1) data that were circularly shifted within game and then concatenated across games, and 2) data where the order of possessions was shuffled within game (preserving possession length) and then concatenated across games. For HMM across-game analyses, we correlated surprise per minute in each game with the number of best fitting HMM-defined states for each game (averaged across subjects), and we assessed significance by scrambling the order of the games. In performing HMM possession-by-possession analyses, we correlated various surprise (and subjective boundary agreement) measures against across-subject HMM state transition agreement, and we assessed chance by circularly shifting the possession order within each game while keeping the order of the games intact. More details on HMM analyses are described above in the *Hidden Markov model (HMM) analyses* section.

For mixed-effects linear and logistic regression models (for pupil area and memory analyses, respectively), we used subject as a random effect and other variables as fixed effects (see Tables S2 and S5 for details). For the pupil area vs surprise time course analysis, we performed mixed-effects linear regression with subject as a random effect and surprise as the fixed effect. Linear and logistic models were created using “lmer” and “glmer” functions in R, respectively. For the neural GLM, we ran subject-specific GLMs and assessed the significance of each factor by taking the beta coefficients from each subject and running a one-sample *t*-test of the betas against zero. For the pupil area, memory, and neural GLMs, we first considered a model containing surprise (along with other regressors); after surprise was found to be significant, we ran follow-up models including belief-consistent and belief-inconsistent surprise as separate factors (Tables S2, S4-5). We avoided including surprise in the same model as belief-consistent and belief-inconsistent surprise to avoid issues related to multicollinearity.

