## Supplementary material for "Behavioral, physiological, and neural signatures of surprise during naturalistic sports viewing": All supplemental figures.

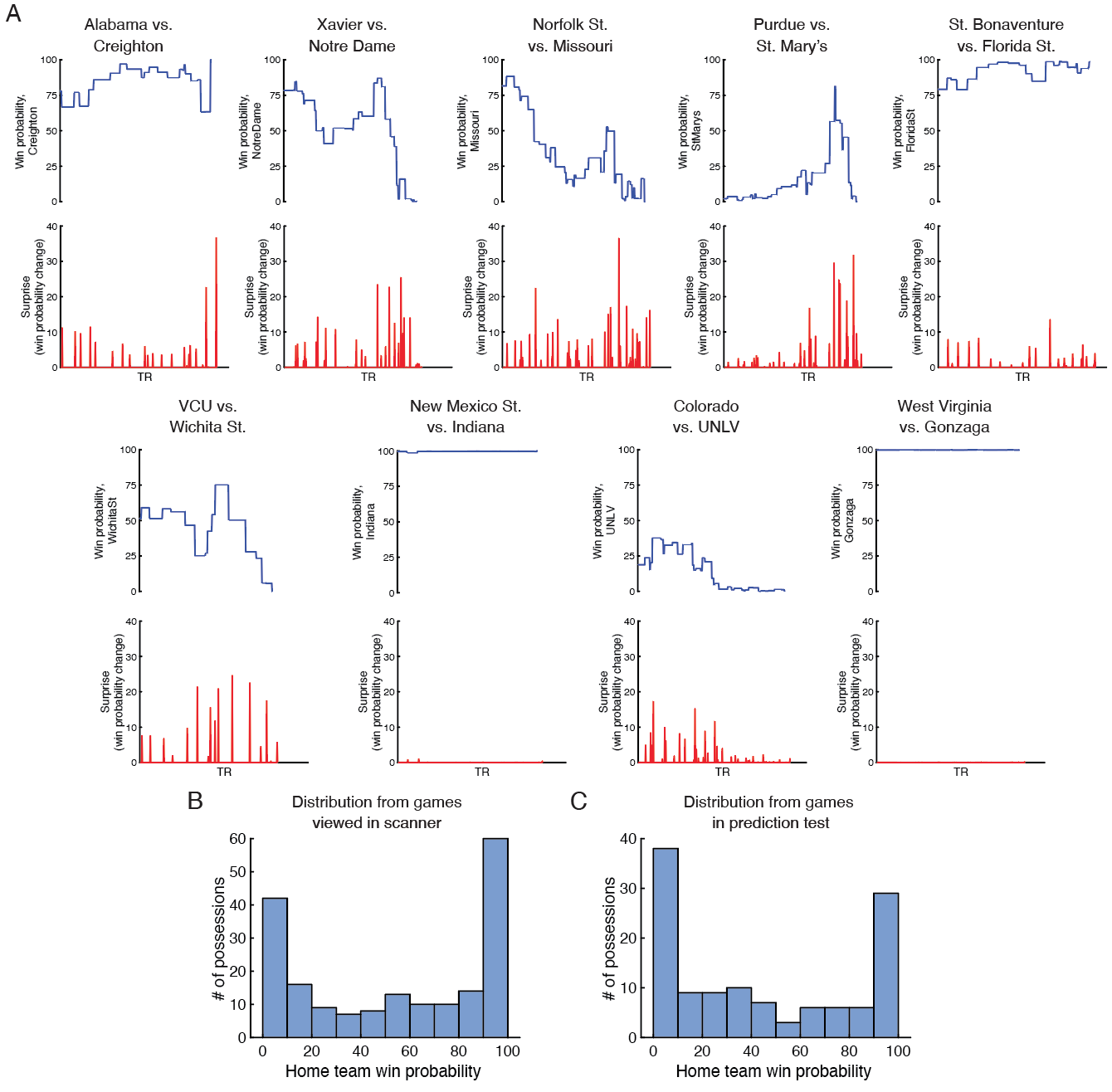

Fig S1. Win probability information, related to Figure 1. a) Win probability and surprise values for all nine games. Similar to Fig 1B. Note that the Indiana and Gonzaga games have near-zero surprise throughout the full clip. b,c) Distribution of win probabilities. Plotted are the number of possessions with win probability values binned every 10% combining all nine games used in the scanner (a) and combining the five games used in the prediction test (b).

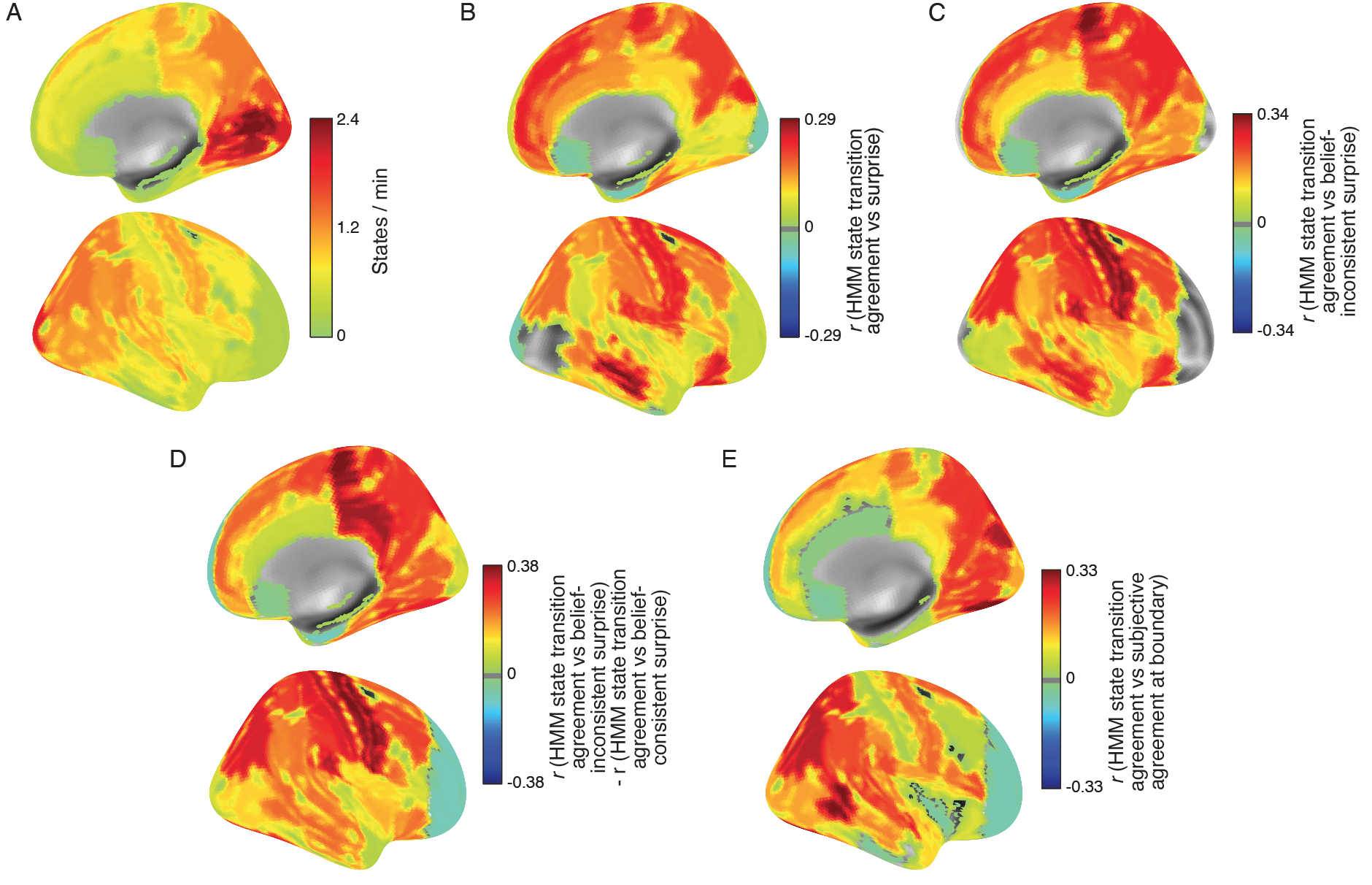

Fig S2. Exploratory analyses applying Hidden Markov models (HMMs) to 48 cortical parcels, related to Fig 3. a) Number of states per minute in each parcel reveals a cortical hierarchy, with the highest number of states in early visual regions and fewer states in higher-level regions. b) Correlation between HMM state transition agreement and surprise at possession boundaries. c) Correlation between HMM state transition agreement and belief-inconsistent surprise at possession boundaries. d) Difference in the correlation between HMM state transition agreement and belief-inconsistent surprise, and the correlation between HMM state transition agreement and belief-consistent surprise. e) Correlation between HMM state transition agreement and subjective boundary agreement at possession boundaries. Data are shown for every parcel and these correlations resemble those done for *a priori* ROIs in Fig 3D. All values, including statistical significance, are in Table S1.

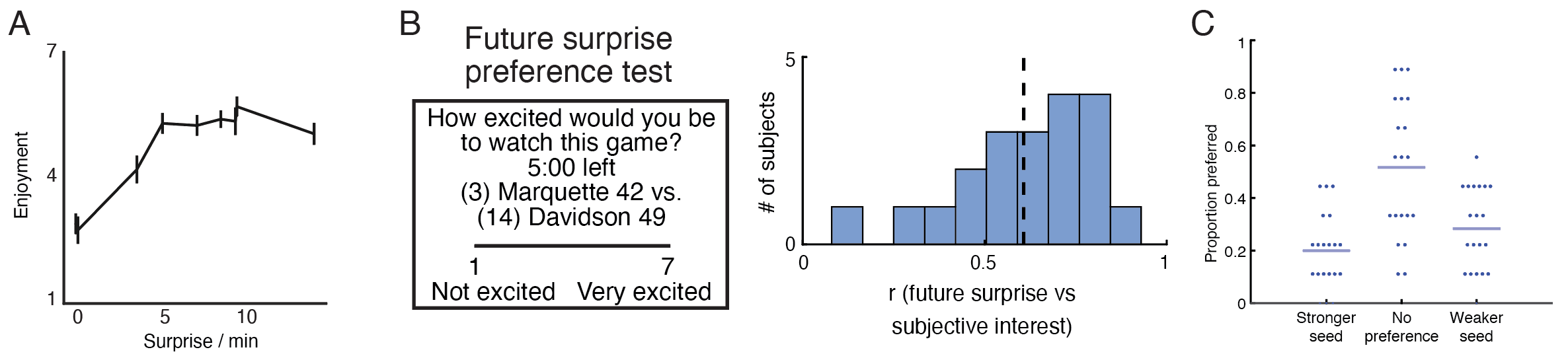

Fig S3: Enjoyment and viewing preferences, related to Fig 5. a) Mean surprise positively predicted game-level enjoyment. Data are represented as mean ± SEM across subjects. b) A survey given at the end of the experiment (left) showed that subjects’ excitement for watching a new set of games strongly correlated with the amount of likely future surprise in those games (as determined by finding similar games in corpus data) (right). c) Proportions of viewed games in which subjects preferred the stronger-seeded team, the weaker-seeded team, or had no preference. These results show that subjects had a preference in approximately half of the games.

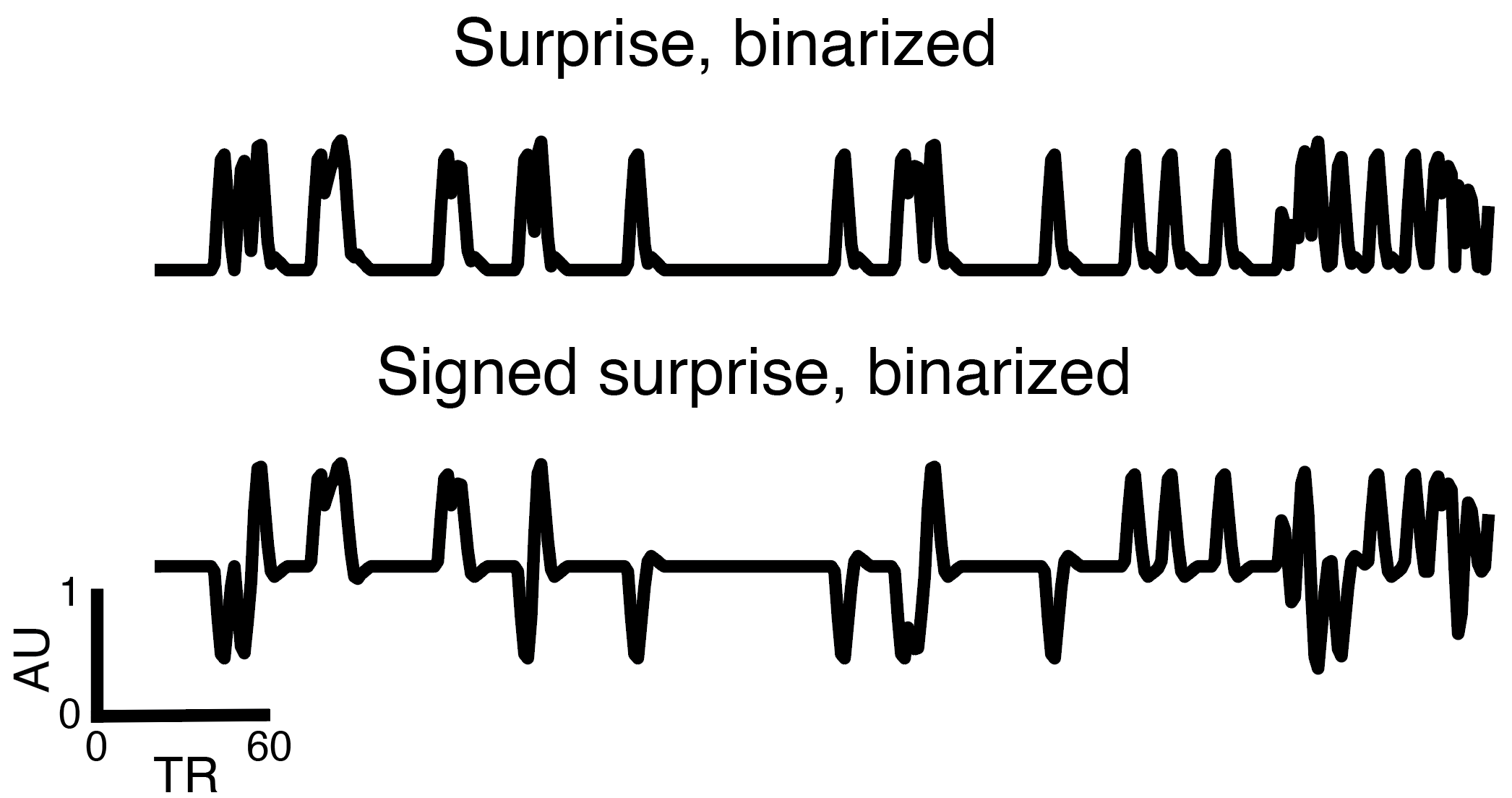

Fig S4. Depictions for one game of binarized “onset” regressors of surprise and signed surprise, related to Fig 5. The goal of including these regressors against the regular versions is to rule out alternative explanations that a neural region may simply respond to any unsigned or signed response in an equal fashion rather than to the actual graded extent of the model-derived surprise.

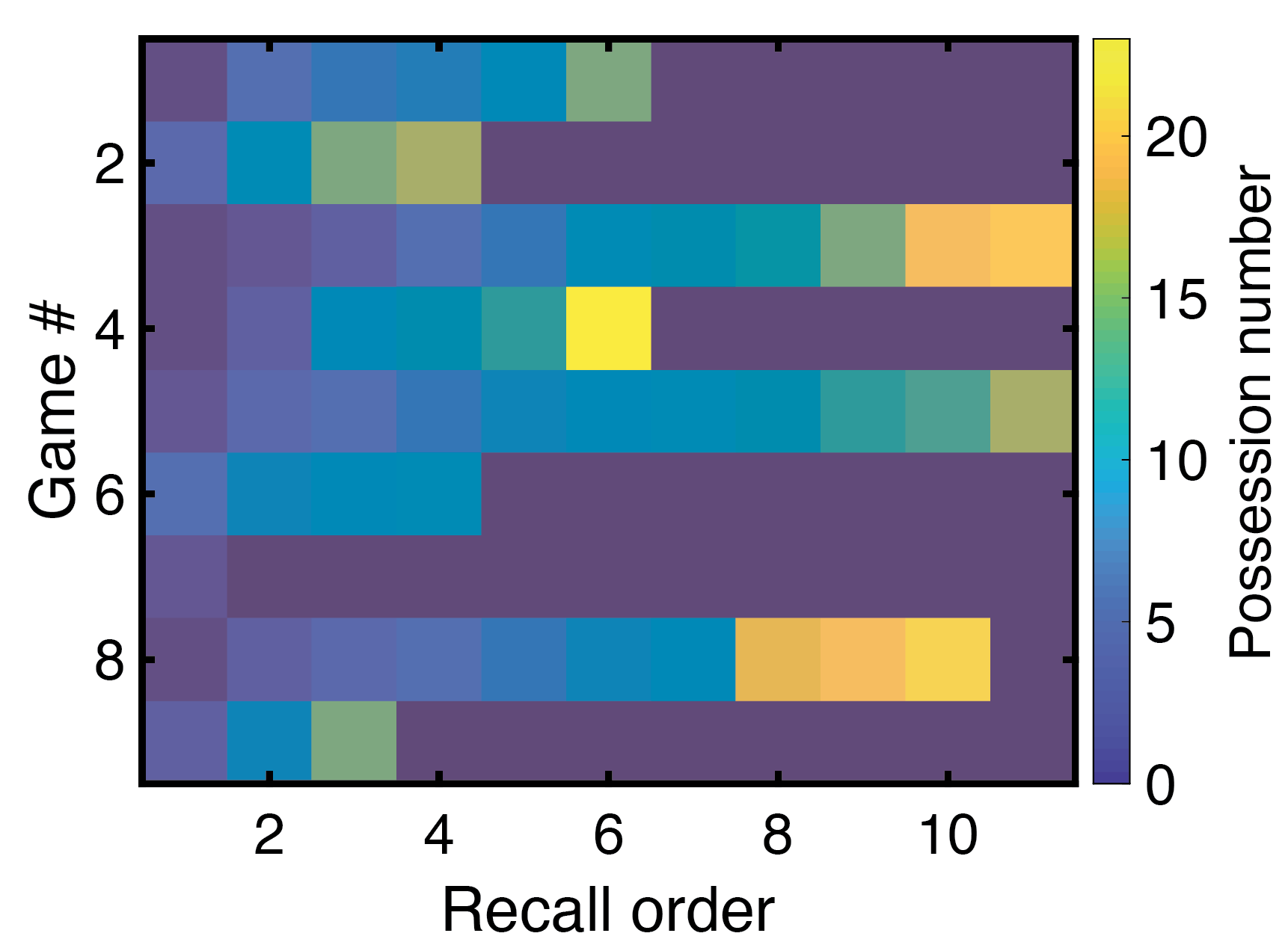

Fig S5. Sample recall transition structure across possessions for one subject, related to Fig 6. The order of recall (x-axis) is shown for all games (y-axis), with the corresponding possession number from the game depicted in color. These results show no backward transitions in recall across possessions.

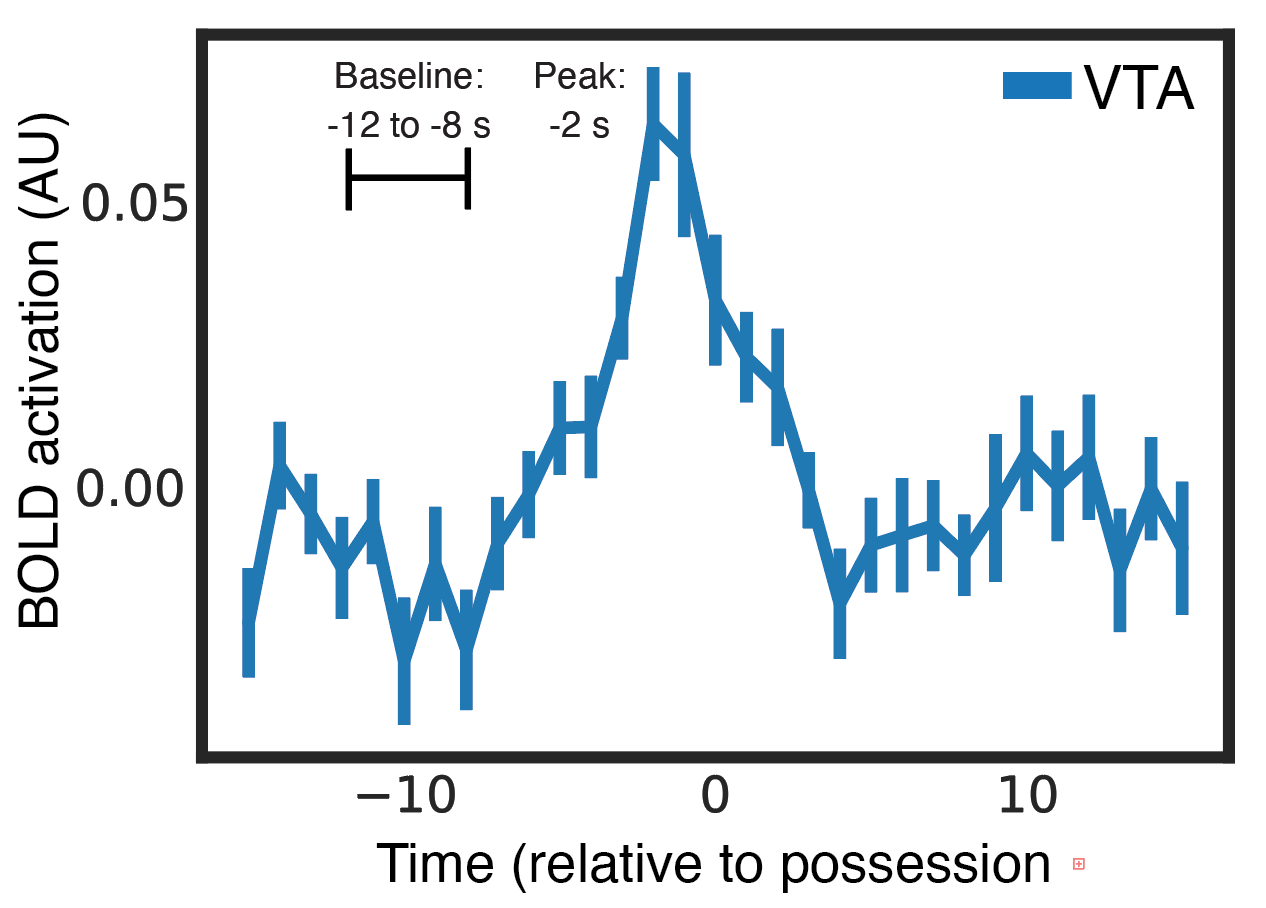

Fig S6. BOLD VTA activation relative to possession boundaries for time-shifted (5 s) data, related to STAR Methods. We used peak activity at -2 s (subtracting a -12 to -8 s baseline) to predict memory on a trial-by-trial basis. Data are represented as mean ± SEM across subjects.

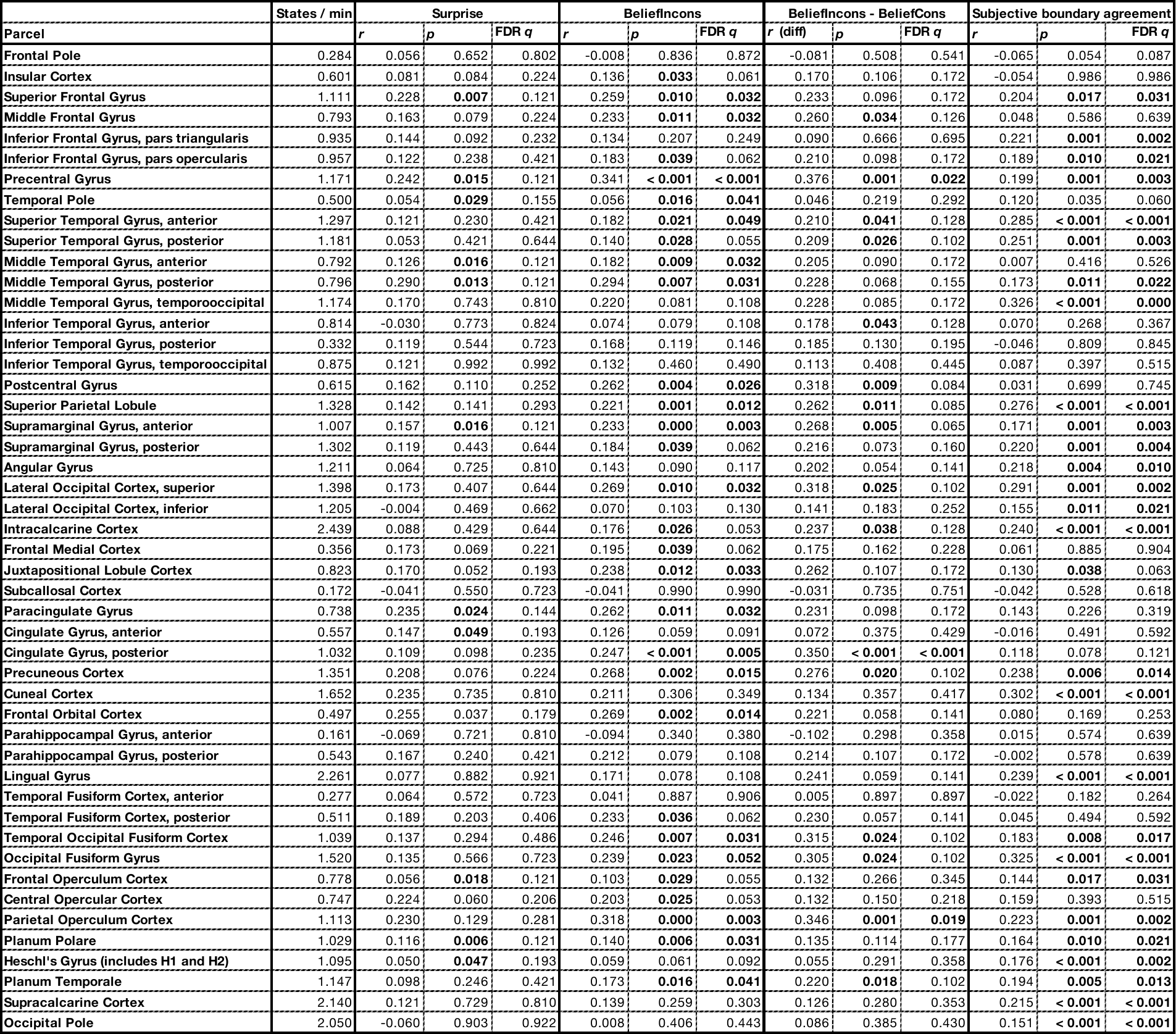

Table S1. Number of HMM states per minute and correlations between HMM state transition agreement at boundaries with various measures of surprise and subjective boundary agreement, related to Fig 3. Uncorrected p values and FDR-corrected q values are provided for each correlation based on permutation tests using within-game circular shifts. BeliefCons = belief-consistent surprise, BeliefIncons = belief-inconsistent surprise. These data are plotted in Fig S2. Values in bold indicate significance at the 0.05 level.

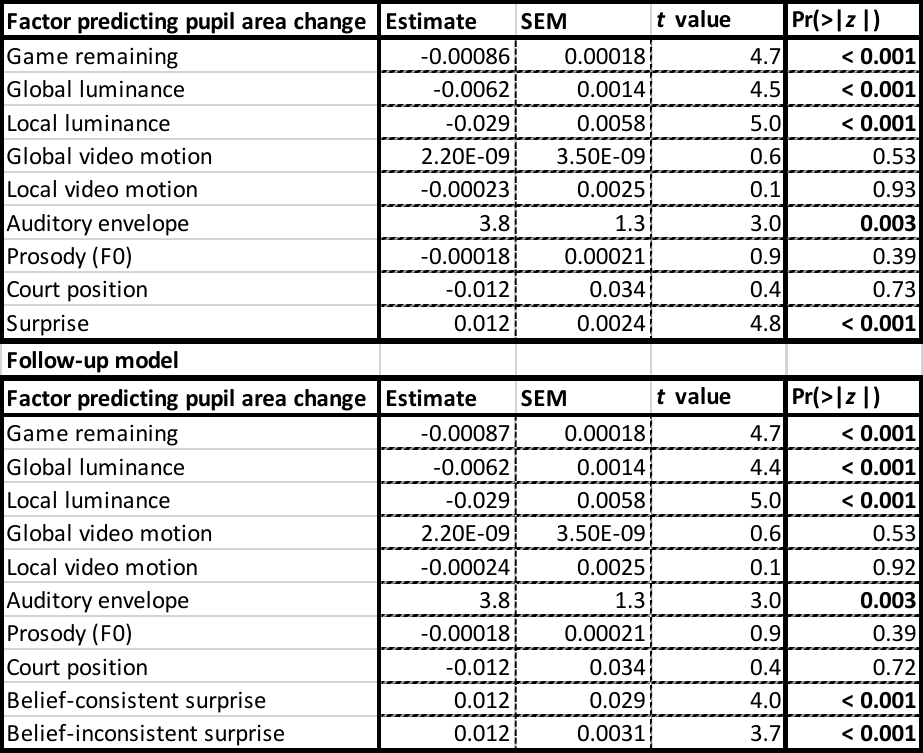

Table S2. Factors predicting pupil area change across the possession boundary in original (top) and follow-up model (bottom), related to Fig 4. The follow-up model was run to dissociate surprise (which was significant in the original model) into belief-consistent and belief-inconsistent surprise. Values in bold indicate significance at the 0.05 level.

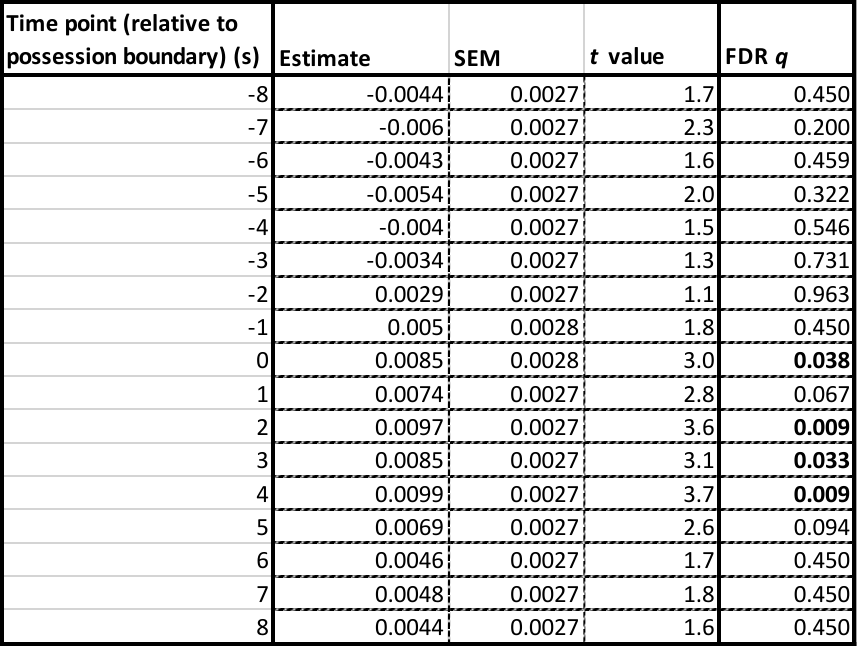

Table S3. Effects of surprise on pupil area in separate models trained at each time point with respect to the possession boundary using surprise as a fixed effect and subject as a random effect, related to Fig 4. Values in bold indicate significance at the 0.05 level.

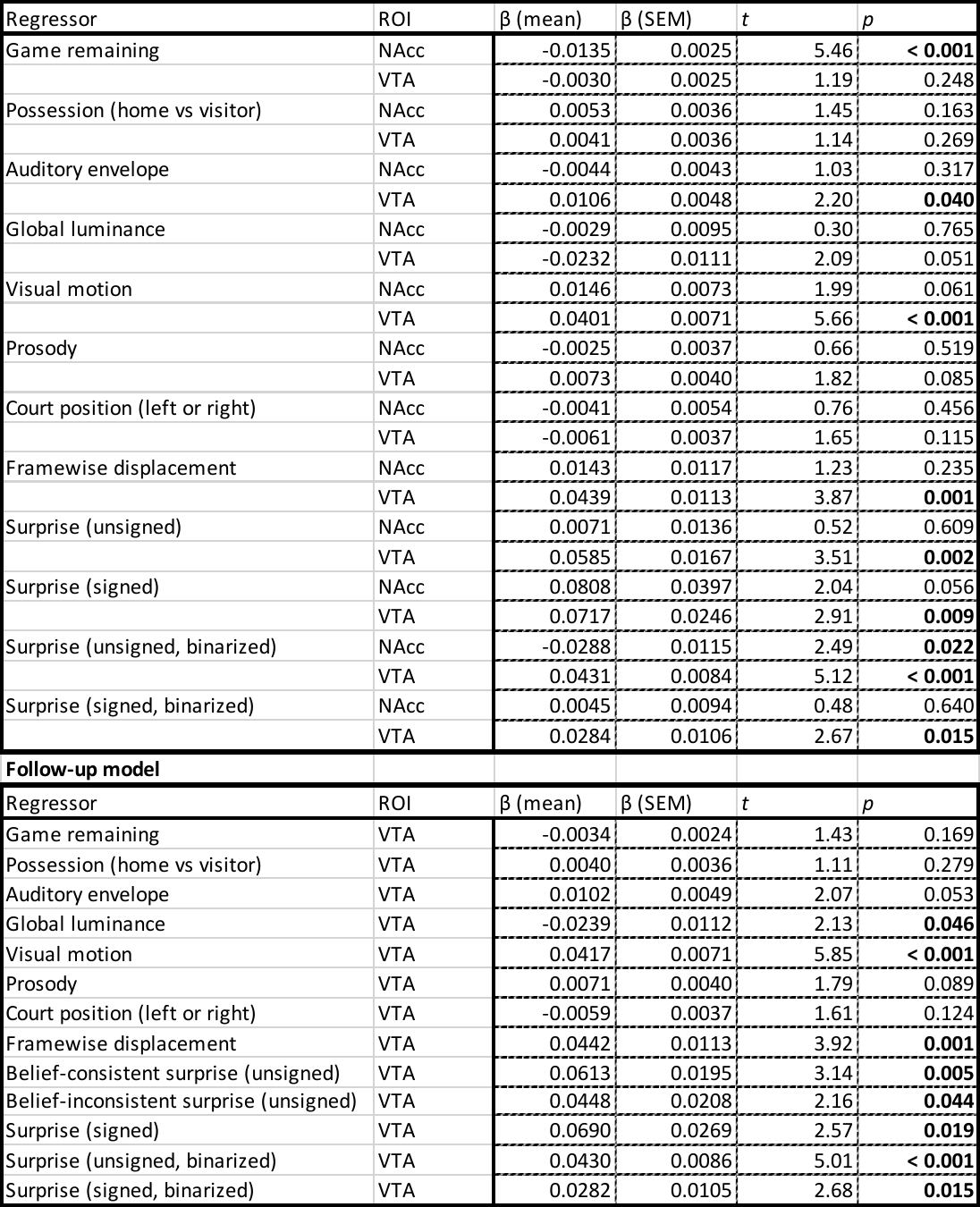

Table S4. Full details of general linear model (GLM) analyses run separately on the nucleus accumbens (NAcc) and ventral tegmental area (VTA), related to Fig 5. Values in bold indicate significance at the 0.05 level.

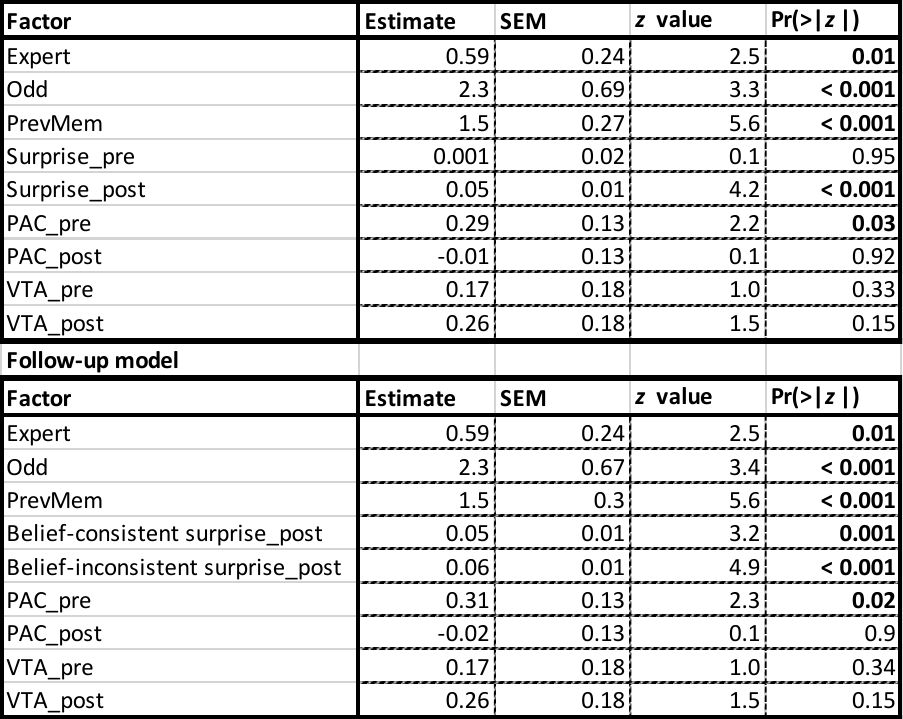

Table S5. Factors predicting memory for possessions, related to Fig 6. Note: Expert = subject expertise, Odd = oddness of possession, PrevMem = previous possession recalled, PAC = pupil area change, VTA = ventral tegmental area activation. “_pre” indicates the value leading into the possession, whereas “_post” indicates the value at the end of the possession. Values in bold indicate significance at the 0.05 level.
